# Estimating cell-type-specific gene co-expression networks from bulk gene expression data with an application to Alzheimer’s disease

**DOI:** 10.1101/2021.12.21.473558

**Authors:** Chang Su, Jingfei Zhang, Hongyu Zhao

## Abstract

Inferring and characterizing gene co-expression networks has led to important insights on the molecular mechanisms of complex diseases. Most co-expression analyses to date have been performed on gene expression data collected from bulk tissues with different cell type compositions across samples. As a result, the co-expression estimates only offer an aggregate view of the underlying gene regulations and can be confounded by heterogeneity in cell type compositions, failing to reveal gene coordination that may be distinct across different cell types. In this paper, we describe a flexible framework for estimating cell-type-specific gene co-expression networks from bulk sample data, without making specific assumptions on the distributions of gene expression profiles in different cell types. We develop a novel sparse least squares estimator, referred to as CSNet, that is efficient to implement and has good theoretical properties. Using CSNet, we analyzed the bulk gene expression data from a cohort study on Alzheimer’s disease and identified previously unknown cell-type-specific co-expressions among Alzheimer’s disease risk genes, suggesting cell-type-specific disease pathology for Alzheimer’s disease.

## 1 Introduction

Gene co-expression networks characterize correlations of gene expression levels across biological samples and co-expressed genes may be regulated by the same transcription factors, functionally related, or involved in the same pathways (Gaiteri et al., 2014). In the past decades, gene co-expression networks have been extensively used to identify functional modules of genes and pathways, which were further associated with disease phenotypes (Wang et al., 2016; Mostafavi et al., 2018; Meng and Mei, 2019; Wang et al., 2021b). For example, a gene co-expression analysis in Zhang et al. (2013) identified TYROBP as a key regulator in an immune related module upregulated in late-onset Alzheimer’s disease, which was recapitulated *in vivo* in mice and is now a new therapeutic target for the disease.

While the literature on gene co-expression networks is readily growing, most co-expression analyses to date have been performed on data collected from bulk tissue samples that aggregate the expression profiles from different cell types. As a result, the estimated co-expression networks only offer an aggregated view of the underlying gene regulations, while gene regulations may differ considerably in different cell types (Heintzman et al., 2009), and the co-expression estimates can be dominated by signals from the more abundant cell types. Moreover, as different bulk samples may have different cell type compositions, the observed co-expressions may be confounded with cell type proportions. For example, consider two genes that are both highly expressed in one cell type but are not co-expressed at the cell type level. Bulk sample data may show that these two genes co-expressed as their expression levels co-vary with the proportion of this specific cell type in the bulk sample. To avoid such confounded results and to gain a more accurate and comprehensive view of the underlying biological processes, a better approach is to estimate cell-type-specific co-expression networks.

Cell-type-specific co-expressions can possibly be estimated from single cell RNA sequencing (RNA-seq) data (Hwang et al., 2018) that measure expression profiles in single cells. However, these data are much more limited in the number of biological samples analyzed (Stower, 2019), have high noises due to low coverage and biological noises (Kiselev et al., 2019), and may be biased due to cell isolation and sequencing protocols (Denisenko et al., 2020). Real data results in Section 4 show that cell-type-specific co-expressions estimated from single cell RNA-seq data are noisy and co-expressions are often only seen in the highly abundant cell types.

Instead of resorting to single cell data for cell-type-specific co-expression analysis, we consider the use of bulk sample data for such analysis in this paper. With the readily available rich collections of bulk gene expression data, there is a great need to develop methods for estimating cell-type-specific co-expressions from bulk samples. There is a recent literature on decomposing bulk gene expression profiles into cell-type-specific profiles (Cobos et al., 2020). Using the bulk data, various methods are available to infer mean expression levels in each cell type (Newman et al., 2019), or to infer cell type proportions (Abbas et al., 2009; Wang et al., 2019; Newman et al., 2019; Tang et al., 2020; Jew et al., 2020; Yang et al., 2021). More recently, methods have been proposed to infer cell-type-specific expressions in each sample, such as CIBERSORTx (Newman et al., 2019) and bMIND (Wang et al., 2021a). For each cell type, these methods offer an indirect way to estimate the co-expressions, by calculating the correlations of estimated expression profiles across samples. However, we show in Sections 3 and 4 that these methods rely on either restrictive assumptions, or high-quality external information that is not readily available in practice.

In our work, we consider a different statistical approach and propose a flexible method to estimate **C**ell-type-**S**pecific gene co-expression **Net**works using bulk gene expression data, and call this method `CSNet`. Specifically, we formulate the problem as estimating the means and covariances of unknown densities from different cell types using data (i.e., bulk samples) generated from a convolution of these densities with varying compositions. Our method `CSNet` does not make specific assumptions on the distributions of expression levels from different cell types, and it overcomes the computational challenge in estimating the covariances in a convolution of densities, especially when the number of genes is large, through a novel least squares approach that is efficient to implement and has good theoretical properties. We further propose a sparse estimator with SCAD penalty in the high dimension regime where the number of genes *p* can far exceed the sample size *n*.

Our real data application focuses on the Alzheimer’s disease, a neurodegenerative disorder that causes progressive and irreversible loss of neurons in the brain (Winblad et al., 2016). It is estimated to affect 5.8 million people in the United States and has become the fifth leading cause of death among Americans over 64 years old (Alzheimer’s Association, 2019). Genetic factors are known to play an important role in Alzheimer’s disease, with an estimated heritability of 58–79% for late-onset Alzheimer’s disease, and large scale genome wide association studies (GWAS) have implicated dozens of regions of the human genome for their relevance for Alzheimer’s disease (Sims et al., 2020). To understand the mechanisms of these disease associated risk genes and the pathology of Alzheimer’s disease, gene co-expression networks have been widely employed (Zhang et al., 2013; Wang et al., 2016; Mostafavi et al., 2018; Meng and Mei, 2019; Wan et al., 2020; Wang et al., 2021b). However, most co-expression analyses focus on correlations between bulk samples, that may be confounded with cell type compositions and only offer an aggregated view of the biological processes in different cell types. Recently, more evidence suggests cell-type-specific pathology of Alzheimer’s disease. For example, neuroinflammation represents a key causal pathway in Alzheimer’s disease and involves primarily glial cells in the brain including microglia and astrocytes (Heneka et al., 2015); myelination is also implicated in the disease, which is mainly contributed by oligodendrocytes (Cai and Xiao, 2016). While such evidence is rapidly increasing, there have been very few cell-type-specific co-expression analyses for Alzheimer’s disease. In our analysis, we focused on the bulk RNA-seq data from the Religious Orders Study and Rush Memory and Aging Project (ROSMAP; Bennett et al., 2018), a clinical-pathologic cohort study of Alzheimer’s disease. Using `CSNet`, we estimated gene co-expression networks for four major cell types in the brain, including excitatory neurons, oligodendrocytes, astrocytes and microglia, on genes with known genetic risk for Alzheimer’s disease, where modules of risk genes that uniquely co-express in astrocytes and microglia were uncovered. Both astrocytes and microglia are cell types that are less abundant (less than 20%), and the co-expressions estimated from single cell RNA-seq data showed no co-expressions in these two cell types [see Figure S6(b)]. We have also considered gene sets that function primarily in specific cell types to validate `CSNet` and added several sensitivity analyses to further validate our results.

The rest of the paper is organized as follows. Section 2 introduces the problem and discusses estimating cell-type-specific co-expressions from bulk samples using `CSNet`. Section 3 reports the simulation results, and Section 4 conducts an analysis of cell-type-specific gene co-expression networks on gene sets with known cell-type-specific functions and an Alzheimer’s disease risk gene set using bulk RNA-seq data from the ROSMAP study. Section 5 investigates theoretical properties of the proposed estimator. Section 6 concludes the paper with a brief discussion.

## 2 Model and Estimation

### 2.1 Problem formulation

Suppose we have expression data ***x***_1_, …, ***x***_*n*_ ∈ ℝ^*p*^ collected from *n* bulk RNA-seq samples across *p* genes. We assume that there are *K* cell types, and the observed bulk level expression is the sum of these *K* cell types written as

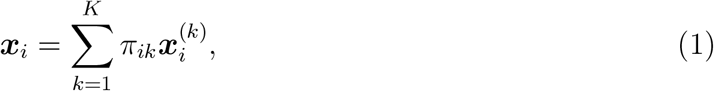

where *π*_*ik*_ and 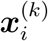 represent the proportion and expression profile of the *k*th cell type in the *i*th sample, respectively. Let 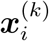 be independent from a multivariate distribution with mean ***μ***^(*k*)^ ∈ ℝ^*p*^ and covariance Σ^(*k*)^ ∈ ℝ^*p×p*^, where ***μ***^(*k*)^ and Σ^(*k*)^ characterize, respectively, the cell-type-specific mean gene expression and co-expression across samples. As the gene regulation mechanisms in functionally distinct cell types are different (Heintzman et al., 2009), we further assume that 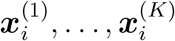 are independent (see discussions in Section 6). Correspondingly, we can write

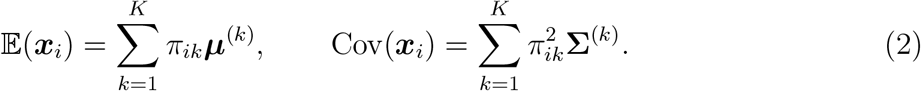

As Σ^(*k*)^ does not relate directly to the strength of gene co-expressions, due to heterogeneity in variances, we further consider correlation matrices in our analysis denoted as 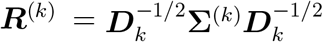, where ***D***_*k*_ is a *p × p* diagonal matrix with the same diagonal as Σ^(*k*)^.

Denote [*m*] = {1, 2, …, *m*} for a positive integer *m*. We shall assume the cell-type proportions *π*_*ik*_’s in (2) are given in the ensuing development, and later demonstrate in experiments that our method is not sensitive to uncertainties in *π*_*ik*_’s; see discussions in Section 6. To infer *π*_*ik*_’s from bulk samples, many methods have been developed that utilize cell type marker genes (i.e., genes that are only highly expressed in one cell type of interest) with expressions profiles gathered from pure cell types (Newman et al., 2015; Li et al., 2016) or single cell RNA-seq data (Wang et al., 2019; Newman et al., 2019; Jew et al., 2020; Dong et al., 2021; Yang et al., 2021). In these methods, the proportions *π*_*ik*_’s are estimated by, for example, nonnegative least squares (Wang et al., 2019) or support vector regression (Newman et al., 2019). Given the bulk samples {***x***_*i*_}_*i*∈ [*n*]_ and cell-type proportions {*π*_*ik*_}_*i*∈[*n*],*k*∈ [*K*]_, our goal is to estimate the cell-type-specific correlations {***R***^(*k*)^}_*k*∈[*K*]_.

### 2.2 Estimating ***R***^(*k*)^ with large *p*

It is easily seen from (1) and (2) that each bulk sample ***x***_*i*_ is from a convolution of *K* distributions. In this case, estimating {Σ^(*k*)^}_*k*∈ [*K*]_ from {***x***_*i*_}_*i*∈ [*n*]_ is very challenging. For example, even in the simple Gaussian case, the loglikelihood function is, up to a constant,

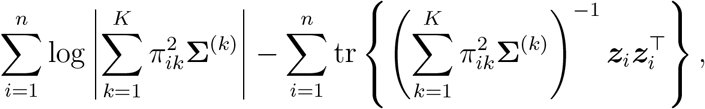

where tr(*·*) denotes the trace of a matrix and ***z***_*i*_ = ***x***_*i*_−𝔼(***x***_*i*_). This loglikelihood is not convex or biconvex with respect to {Σ^(*k*)^}_*k*∈[*K*]_, and cannot be directly optimized using existing iterative algorithmic solutions such as EM and coordinate descent. To our knowledge, there are no existing methods that can effectively estimate the covariances in a convolution of densities, especially when *p>n* as in our problem.

To tackle this challenge, we propose a novel moment-based approach that is efficient to implement and flexible, in that it does not assume the distributions from the *K* cell types to be known or of the same type. The proposed approach, named `CSNet`, first estimates ***R***^(*k*)^ efficiently in an element-wise fashion and then applies a thresholding step, in the case of a large *p*, to give a sparse estimator. Next, we introduce the `CSNet` estimator.

Letting 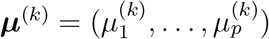 and 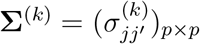, (1) and (2) together imply

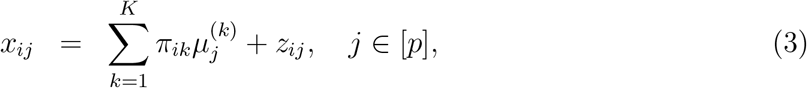

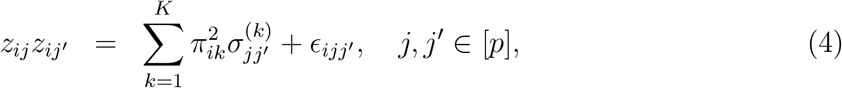

where 𝔼(*z*_*ij*_) = 0 and 𝔼(*ϵ*_*ijj*′_) = 0. This formulation facilitates an efficient least squares estimation procedure to be detailed in the next paragraph. Note that (3)-(4) hold generally without parametric assumptions on the distributions from the *K* cell types.

Denote ***y***_*j*_ = (*x*_1*j*_, …, *x*_*nj*_) and ***D*** = (*π*_*ik*_)_*n×K*_. Equation (3) entails estimation of the cell-type-specific mean ***μ***^(*k*)^ via

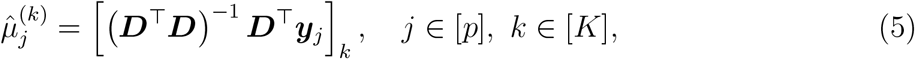

where [***x***]_*k*_ is the *k*th entry in ***x*** ∈ ℝ^*K*^. Let 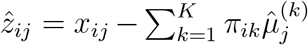 and 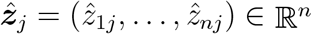. Denoting 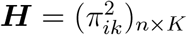, equation (4) entails estimation of the cell-type-specific covariance 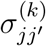 via

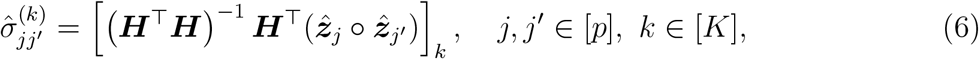

where ° denotes element-wise product. Then, the cell-type-specific correlation 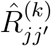 is estimated as

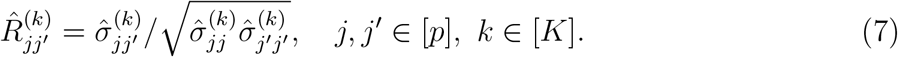

In Section A3.2, we show that element-wisely 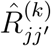 is consistent with a 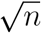-convergence rate under certain regularity conditions.

Though each entry in the correlation matrix estimated in (7) has a 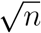-convergence rate, the accumulated errors across *O*(*p*^2^) entries in 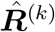 can be excessive, especially as the number of genes *p* often far exceeds the sample size *n* in co-expression network analysis, which can negatively impact downstream analyses such as ranking, principal component analysis or clustering. This same challenge also arises in estimating large sample covariance (Bickel and Levina, 2008a,b; Rothman et al., 2008, 2009) and correlation matrices (El Karoui, 2008; Jiang, 2013). To facilitate estimability and interpretability, we assume that Σ^(*k*)^ (or equivalently ***R***^(*k*)^) is approximately sparse for all *k*; see the definition of approximate sparsity in Assumption 2. Sparsity is plausible in our data problem, as gene co-expressions are expected to be sparse when *p* is large (Zhang and Horvath, 2005).

Based on 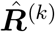, the proposed `CSNet` estimator computes sparse cell-type-specific correlation estimates via thresholding. Specifically, `CSNet` applies an element-wise SCAD (Fan and Li, 2001) thresholding operator to 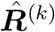, written as 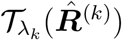, with the (*j, j*′)th thresholded entry calculated as

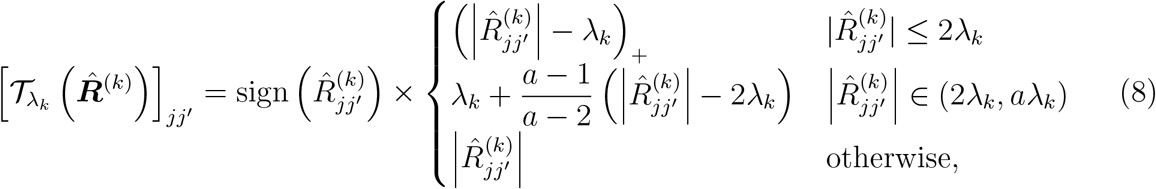

where *λ*_*k*_ is a tuning parameter and we discuss its selection in Section 2.3. As has been well established in the sparse covariance estimation literature (e.g. Bickel and Levina, 2008a,b; Rothman et al., 2008, 2009), the thresholding procedure is easy to implement and enjoys good theoretical properties. Moreover, the SCAD thresholding has been found to give better numerical performances when compared to soft or hard thresholding (Rothman et al., 2009). We set *a* = 3.7 in our experiments as recommended by Fan and Li (2001). The thresholds *λ*_*k*_’s may differ across different cell types to accommodate varying sparsity among different cell types. However, the *λ*_*k*_’s can be selected separately (without a joint tuning) in our tuning procedure (see Section 2.3), which is an attractive computational property of our procedure. In Section 5, we show the convergence rates of `CSNet`, i.e., 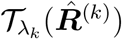, in spectral and Frobenius norms, and establish its selection consistency.

As with all thresholding approaches, 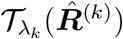 is not guaranteed to be positive definite, though it is asymptotically positive definite as ensured by Theorems 5.1-5.2; see more discussions in Section 6 on considerations for positive definiteness. To ensure the finite sample validity of correlation estimates, we threshold the correlation estimates to be within [-1,1] in our experiments.

### 2.3 Time complexity and parameter tuning

We first discuss the time complexity of solving (6) for all entries in covariance matrices. Though (6) is computed element-wisely, the matrix (***H***^T^***H*)**^***−*** 1^ ***H***^T^ ∈ ℝ^*K×n*^ is common and only needs to be calculated once. Hence, entries in Σ_1_, …, Σ_*K*_ can be estimated efficiently via (***H***^T^***H*)**^***−*** 1^ ***H***^T^***Y***, where ***Y*** is an *n × p*^2^ matrix with the *jj*′th column set to 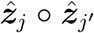. Correspondingly, the time complexity of estimating Σ_1_, …, Σ_*K*_ is *O*(*Kn*^2^*p*^2^), while that of a naive sample covariance estimation is *O*(*np*^2^). As the number of cell types *K* is usually small, the proposed element-wise estimation is computationally feasible when *n*, the number of bulk RNA-seq samples, is moderate.

Next, we discuss parameter tuning. In our procedure, the tuning parameters *λ*_*k*_’s are selected using cross validation or, if available, an independent validation data set. Here, we introduce the cross validation procedure and note that selection with a validation data set can be carried out similarly. We randomly split the data into two equal-sized pieces and estimate for each piece, the cell-type-specific correlation matrices as in (7). We denote the estimated correlation matrices from these two data splits as 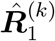 and 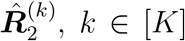, respectively. For each *k*, the tuning parameter *λ*_*k*_ is selected by minimizing 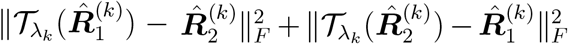 among a set of working values, where *‖ ·‖*_*F*_ denotes Frobenius norm. We consider two equal sized data splits as sufficient samples are needed to estimate 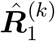 and 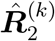 well. This procedure is similar to what was proposed in Bickel and Levina (2008a), where the theoretical justification was provided, and it is found to give a good performance in our numerical experiments. An attractive feature of our proposed tuning procedure is that *λ*_*k*_’s are selected separately without the need of a joint tuning, further reducing the computational cost.

## 3 Simulation Studies

We first investigate the finite sample performance of `CSNet` and compare it with two existing methods, and then conduct a sensitive analysis to examine the performance of `CSNet` when provided cell type proportions are inaccurate. A notable contribution from our simulation studies is a new and efficient method for simulating from multivariate negative binomial distributions with large and structured covariances, presented in Algorithm 1 in the supplement, which can be of independent interest.

### 3.1 Comparing CSNet with other methods

In this section, we consider [1] the proposed `CSNet` estimator calculated as in (8), [2] the cell-type-specific estimator in (7) without sparsity, referred to as `d-CSNet`, [3] the sparse estimator with SCAD thresholding in Rothman et al. (2009) for bulk samples (i.e., not cell-type-specific), referred to as `Bulk`, and [4] the estimator computed from the sample level cell-type-specific expressions estimated by `bMIND` (Wang et al., 2021a). The method `bMIND` considers a Bayesian mixed effects model that constructs priors using single cell data and estimates cell-type-specific expressions in each sample with posterior means. In our experiments, `bMIND` is evaluated with non-informative priors to be comparable with the other methods, which do not depend on prior knowledge. We also consider informative priors for `bMIND` in Tables S1 and S3 in the supplement and the results remain similar. We chose to not compare with CIBERSORTx high-resolution expression purification (Newman et al., 2019) as it is applicable only when there are both case and control samples in the data. Both `CSNet` and `d-CSNet` are computed using the procedure in Section 2.2 with nonnegative least squares to estimate the unknown ***μ***^(*k*)^’s.

We simulate *n* bulk samples of dimension *p* following 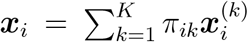 in (1) with *K* = 2 cell types and *π*_*i*1_’s i.i.d from Beta(2, 1). Correspondingly, cell type 1 (*m* = 2*/*3) is on average twice as abundant as cell type 2 (*m* = 1*/*3), where *m* denotes the average cell type proportions. We simulate 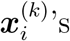 from multivariate negative binomial distributions, resembling read counts from bulk RNA-seq data (Love et al., 2014), with mean ***μ***^(*k*)^’s and covariance matrices Σ^(*k*)^’s specified as follows. The *p* genes are divided into three equal-sized sets, denoted as *V*_1_, *V*_2_ and *V*_3_; genes in *V*_1_ and *V*_2_ are set to co-express in cell types 1 and 2, respectively, while all other correlations are set to zero (see the left panel in Figure 1). For the co-expressed genes in *V*_1_ or *V*_2_, two types of structures are considered, including an MA(1) structure with *ρ*_*jj*′_ = 0.39*×*1_|*j−j*′|=1_ and an AR(1)-type structure with *ρ*_*jj*′_ = 0.70*×*0.9^|*j− j*′ *−* 1|^, for *j ≠ j*′ ∈ *V*_1_ or *V*_2_. We set log 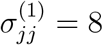 for all *j*, log 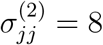 for *j* ∈ *V*_1_, *V*_2_ and log 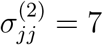 for *j* ∈ *V*_3_ with sequencing depth set to *S* = 6*×*10^7^ in Algorithm 1 to mimic highly-expressed protein-coding genes in real sequencing data. The mean ***μ***^(*k*)^ is set to be a function of Σ^(*k*)^ (see details in Algorithm1), consistent with the observation in real data that higher expression levels are often associated with larger variances. The tuning parameters for `CSNet` and `Bulk` are selected following Section 2.3 and the suggested procedure in Rothman et al. (2009), respectively, both using a validation data set with 150 independent samples. We consider network sizes *p* = 100, 200 and sample sizes *n* = 150, 600.

**Figure 1:**
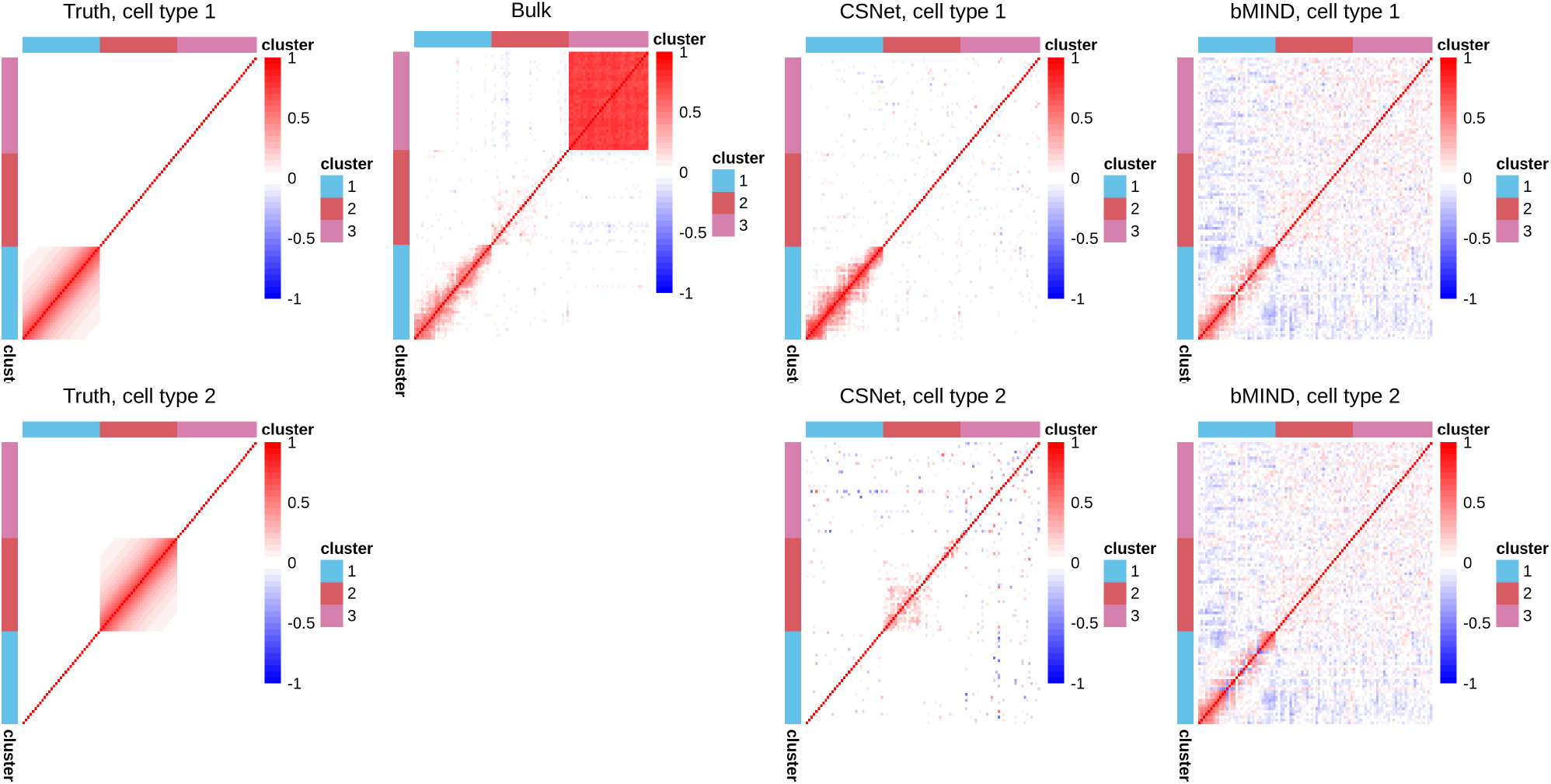
Correlation matrices in one data replicate with *p* = 100 and *n* = 150, under a negative binomial distribution with an AR(1)-type correlation structure in *V*_1_ or *V*_2_ (See Section 3). From left to right: true correlation matrices, estimates from `Bulk`, `CSNet` and `bMIND`. Note the `Bulk` estimate is not cell-type-specific.

To evaluate the estimation accuracy, we report the estimation errors in Frobenius norm 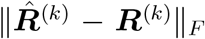 and operator norm 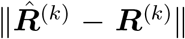, where 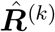, with a slight overuse of notation, denotes the estimate of ***R***^(*k*)^ obtained by various methods. Also reported are the true positive rate (TPR) and false positive rate (FPR), which evaluate the selection accuracy of nonzero entries in ***R***^(*k*)^’s. The TPR and FPR are only reported for sparse estimators `Bulk` and `CSNet`. Table 1 reports the average criteria under the MA(1) model for four methods, with standard deviations in the parentheses, over 200 data replications. The results for AR(1)-type are similar and relegated to Table S2 in the supplement due to space limitation. It shows that `CSNet` achieves the best performance in terms of both estimation accuracy and selection accuracy. In the supplement, we demonstrate in Tables S1 and S3 that even with informative priors derived from simulated cell-type-specific data, `CSNet` still performs better than `bMIND`.

**Table 1:**
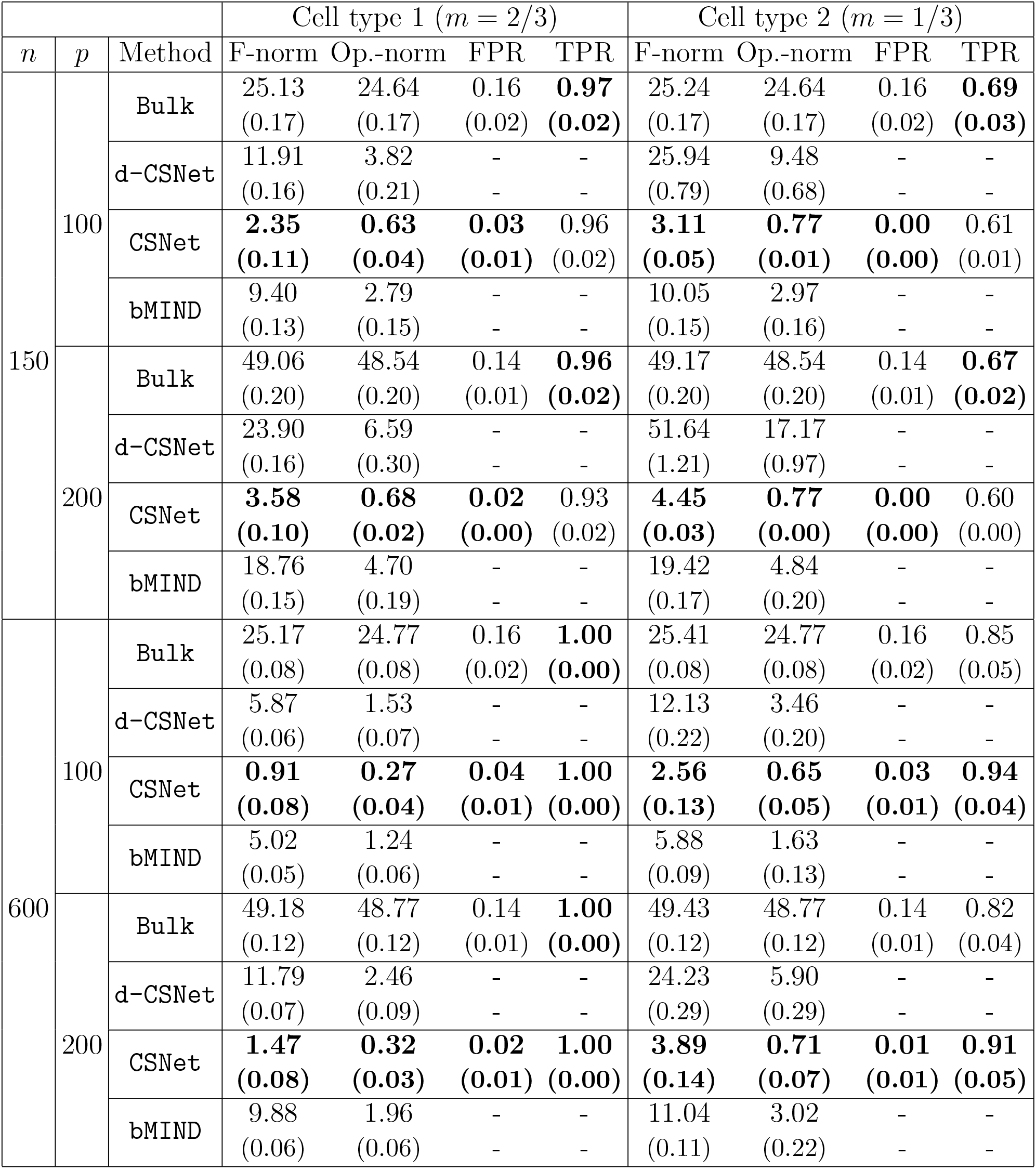
Evaluation criteria with varying sample size *n* and network size *p* under a negative binomial distribution with an MA(1) correlation structure in *V*_1_ or *V*_2_ (See Section 3). The four methods under comparison are `Bulk`, `d-CSNet`, `CSNet` and `bMIND`. We use F-norm to denote the Frobenius norm and Op.-norm to denote the operator norm. Marked in boldface are those achieving the best evaluation criteria in each setting.

| | | | Cell type 1 ($m = 2/3$) | | | | Cell type 2 ($m = 1/3$) | | | |
| --- | --- | --- | --- | --- | --- | --- | --- | --- | --- | --- |
| $n$ | $p$ | Method | F-norm | Op.-norm | FPR | TPR | F-norm | Op.-norm | FPR | TPR |
| 150 | 100 | Bulk | 25.13<br>(0.17) | 24.64<br>(0.17) | 0.16<br>(0.02) | **0.97**<br>(**0.02**) | 25.24<br>(0.17) | 24.64<br>(0.17) | 0.16<br>(0.02) | **0.69**<br>(**0.03**) |
|  |  | d-CSNet | 11.91<br>(0.16) | 3.82<br>(0.21) | -<br>- | -<br>- | 25.94<br>(0.79) | 9.48<br>(0.68) | -<br>- | -<br>- |
|  |  | CSNet | **2.35**<br>(**0.11**) | **0.63**<br>(**0.04**) | **0.03**<br>(**0.01**) | 0.96<br>(0.02) | **3.11**<br>(**0.05**) | **0.77**<br>(**0.01**) | **0.00**<br>(**0.00**) | 0.61<br>(0.01) |
|  |  | bMIND | 9.40<br>(0.13) | 2.79<br>(0.15) | -<br>- | -<br>- | 10.05<br>(0.15) | 2.97<br>(0.16) | -<br>- | -<br>- |
|  | 200 | Bulk | 49.06<br>(0.20) | 48.54<br>(0.20) | 0.14<br>(0.01) | **0.96**<br>(**0.02**) | 49.17<br>(0.20) | 48.54<br>(0.20) | 0.14<br>(0.01) | **0.67**<br>(**0.02**) |
|  |  | d-CSNet | 23.90<br>(0.16) | 6.59<br>(0.30) | -<br>- | -<br>- | 51.64<br>(1.21) | 17.17<br>(0.97) | -<br>- | -<br>- |
|  |  | CSNet | **3.58**<br>(**0.10**) | **0.68**<br>(**0.02**) | **0.02**<br>(**0.00**) | 0.93<br>(0.02) | **4.45**<br>(**0.03**) | **0.77**<br>(**0.00**) | **0.00**<br>(**0.00**) | 0.60<br>(0.00) |
|  |  | bMIND | 18.76<br>(0.15) | 4.70<br>(0.19) | -<br>- | -<br>- | 19.42<br>(0.17) | 4.84<br>(0.20) | -<br>- | -<br>- |
| 600 | 100 | Bulk | 25.17<br>(0.08) | 24.77<br>(0.08) | 0.16<br>(0.02) | **1.00**<br>(**0.00**) | 25.41<br>(0.08) | 24.77<br>(0.08) | 0.16<br>(0.02) | 0.85<br>(0.05) |
|  |  | d-CSNet | 5.87<br>(0.06) | 1.53<br>(0.07) | -<br>- | -<br>- | 12.13<br>(0.22) | 3.46<br>(0.20) | -<br>- | -<br>- |
|  |  | CSNet | **0.91**<br>(**0.08**) | **0.27**<br>(**0.04**) | **0.04**<br>(**0.01**) | **1.00**<br>(**0.00**) | **2.56**<br>(**0.13**) | **0.65**<br>(**0.05**) | **0.03**<br>(**0.01**) | **0.94**<br>(**0.04**) |
|  |  | bMIND | 5.02<br>(0.05) | 1.24<br>(0.06) | -<br>- | -<br>- | 5.88<br>(0.09) | 1.63<br>(0.13) | -<br>- | -<br>- |
|  | 200 | Bulk | 49.18<br>(0.12) | 48.77<br>(0.12) | 0.14<br>(0.01) | **1.00**<br>(**0.00**) | 49.43<br>(0.12) | 48.77<br>(0.12) | 0.14<br>(0.01) | 0.82<br>(0.04) |
|  |  | d-CSNet | 11.79<br>(0.07) | 2.46<br>(0.09) | -<br>- | -<br>- | 24.23<br>(0.29) | 5.90<br>(0.29) | -<br>- | -<br>- |
|  |  | CSNet | **1.47**<br>(**0.08**) | **0.32**<br>(**0.03**) | **0.02**<br>(**0.01**) | **1.00**<br>(**0.00**) | **3.89**<br>(**0.14**) | **0.71**<br>(**0.07**) | **0.01**<br>(**0.01**) | **0.91**<br>(**0.05**) |
|  |  | bMIND | 9.88<br>(0.06) | 1.96<br>(0.06) | -<br>- | -<br>- | 11.04<br>(0.11) | 3.02<br>(0.22) | -<br>- | -<br>- |

To better visualize the estimates, Figure 1 plots heat maps of the true cell-type-specific co-expression matrices and estimates from `Bulk`, `CSNet` and `bMIND`. The estimates from `d-CSNet` show similar patterns as `CSNet` plus some additional noises, and are omitted due to space limitation. From Figure 1, it is clearly seen that both `Bulk` and `bMIND` give a less accurate view of the true co-expressions. Specifically, `Bulk` estimates high co-expressions in *V*_3_ while genes in *V*_3_ are not co-coexpressed in either cell type. True co-expression patterns in *V*_2_ from cell type 2 are also notably attenuated in `Bulk`. Moreover, `bMIND` does not perform well for cell type 2, the less abundent cell type. It is seen that the true co-expressions in *V*_2_ are not identified while co-expressions specific to cell type 1 are incorrectly inferred. In comparison, `CSNet` was able to identify the true co-expression patterns in both cell types.

Finally, to evaluate the threshold selection procedure in Section 2.3, we plot the ROC curves that plot the TPR against the FPR across a fine grid of thresholding parameters for `Bulk`, `CSNet` and `bMIND`. Though `bMIND` is not sparse, we applied the SCAD thresholding operator to the `bMIND` estimates to examine its performance. The thresholds selected by our proposed procedure in Section 2.3 are marked on the curves for `CSNet`. The ROC curves in Figures 2 show that `CSNet` achieves the best performance and the selected thresholds generally strike a reasonable balance between TPR and FPR. As shown in Table 1 and Figures 1-2, the improvement of `CSNet` over others is the most notable for the less abundant cell type, and this demonstrates the efficacy of our proposed method for cell types whose signals are attenuated in bulk samples.

**Figure 2:**
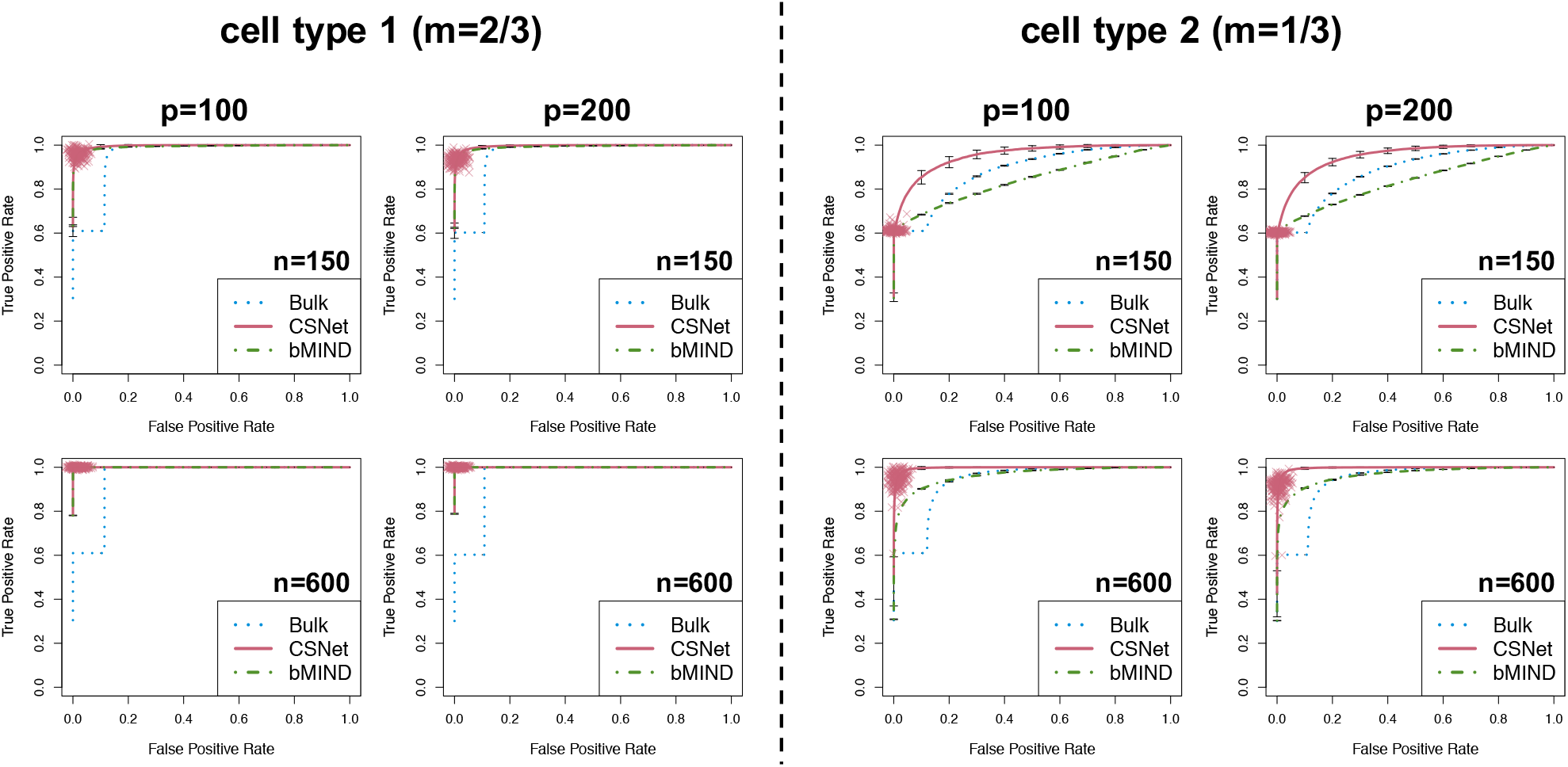
ROC curves with varying sample size *n* and network size *p* under a negative binomial distribution with an MA(1) correlation structure in *V*_1_ or *V*_2_ (See section 3). The three methods under comparison are `Bulk`, `CSNet` and `bMIND`. The thresholds selected by our proposed procedure in Section 2.3 are marked on the curves (in red crosses).

### 3.2 Sensitivity analysis of CSNet

In this section, we conduct a sensitivity analysis to examine the performance of our method when the cell type proportions *π*_*ik*_’s used in `CSNet` are inaccurate. We consider the same simulation setting as in Table 1 with *n* = 150, *p* = 100 and *K* = 2. For *i* ∈ [*n*], let 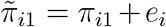 (constrained to be within [0, 1]), where *e*_*i*_’s are independent from *k × N* (0, 0.04) and *k* is set to {0, 0.25, 0.5, 1}. When *k* = 0, 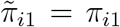 and the cell type proportions are accurate with no errors. The estimation is then carried out following the procedure in Section 2.2 with inaccurate cell type proportions 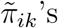.

Table 2 reports the evaluation criteria for estimating cell-type-specific correlation matrices ***R***^(*k*)^’s with `CSNet` under various noise levels. It is seen that under this inaccurate cell type proportion setting, our method still performs reasonably well.

**Table 2:** Sensitivity analysis of `CSNet` with *n* = 150 and *p* = 100, under the same setting as Table 1. Noisy cell type proportions were set to *π*_*i*1_ + *k × N* (0, 0.04), and then thresholded to be within [0, 1].

| | $m = 2/3$ | | | | $m = 1/3$ | | | |
| --- | --- | --- | --- | --- | --- | --- | --- | --- |
|  | F-norm | Op.-norm | FPR | TPR | F-norm | Op.-norm | FPR | TPR |
| $\kappa = 0$ | 2.35<br>(0.11) | 0.63<br>(0.04) | 0.03<br>(0.01) | 0.96<br>(0.02) | 3.11<br>(0.05) | 0.77<br>(0.01) | 0.00<br>(0.00) | 0.61<br>(0.01) |
| $\kappa = 0.25$ | 2.38<br>(0.11) | 0.63<br>(0.04) | 0.03<br>(0.01) | 0.95<br>(0.02) | 3.11<br>(0.05) | 0.77<br>(0.01) | 0.00<br>(0.00) | 0.61<br>(0.01) |
| $\kappa = 0.5$ | 2.72<br>(0.10) | 0.68<br>(0.03) | 0.03<br>(0.01) | 0.89<br>(0.04) | 3.11<br>(0.04) | 0.77<br>(0.00) | 0.00<br>(0.00) | 0.61<br>(0.01) |
| $\kappa = 1$ | 3.09<br>(0.01) | 0.77<br>(0.00) | 0.00<br>(0.00) | 0.62<br>(0.01) | 3.10<br>(0.02) | 0.77<br>(0.00) | 0.00<br>(0.01) | 0.61<br>(0.00) |

## 4 Cell-type-specific Co-expressions of Different Gene Sets for an Alzheimer’s disease Cohort

We focus on estimating cell-type-specific co-expressions using bulk RNA-seq data from the Religious Orders Study and Rush Memory and Aging Project (ROSMAP) (Bennett et al., 2018), a clinical-pathologic cohort study of Alzheimer’s disease. In the ROSMAP study, post-mortem brain samples from *n* = 541 subjects were collected from the grey matter of dorsolateral prefrontal cortex (DLPFC), a brain region heavily implicated in Alzheimer’s disease pathology. Expression unit FPKM (Trapnell et al., 2010) was used to quantify gene expressions, and no notable batch effects were observed from these samples (see Figure S3). The cell type proportions for *K* = 8 cell types were estimated using CIBERSORTx (Newman et al., 2019) with the signature matrix built from the single-nucleus RNA-seq data (Mathys et al., 2019) collected on the same brain region in a subset of 48 samples.

In the ensuing analysis, we focus on four most abundant cell types: excitatory neurons (Ex), oligodendrocytes (Oli), astrocytes (Ast) and microglia (Mic). The average proportions for these four cell types are 0.48, 0.21, 0.18 and 0.08, respectively. We applied `CSNet` as defined in (8), with the tuning parameter selected using the cross validation procedure discussed in Section 2.3. Additionally, we compared `CSNet` to two alternative approaches, one through `bMIND`, the best performing alternative in Section 3, and one using single cell data (Mathys et al., 2019), respectively. For `bMIND` estimates, we followed Wang et al. (2021a) to infer priors from single cell data and supplied these priors to estimate cell-type-specific expressions in each sample; see details in Section A4; correlation estimates were then computed using the estimated cell-type-specific expressions across different samples. To estimate cell-type-specific co-expressions from single cell data, we first calculated cell-type-specific expressions in each sample. Specifically, the expression profile of gene *j* in cell type *k* for sample *i* was calculated by first summing over the UMI counts of gene *j* from all cells of cell type *k* in sample *i*, and then normalized by the total number of UMI counts in cell type *k* from sample *i*. These cell-type-specific expressions calculated for different samples were then used to estimate the co-expression (i.e., correlation matrix) in each cell type, and the correlation matrices were further thresholded following the procedure in Rothman et al. (2009) with the SCAD penalty. For all methods, we visualized the estimated co-expressions using heat maps, with genes ordered into clusters (or modules) identified by WGCNA (Langfelder and Horvath, 2008), a gene clustering method, applied to bulk samples.

### 4.1 Gene sets with known cell-type-specific functions

The gene co-expressions estimated from different methods were compared on a few sets of genes. We first considered three sets of genes obtained from Gene Ontology (GO) (Ashburner et al., 2000; Consortium, 2021) including the *excitatory synapse* genes (GO:0060076, *p* = 46), *myelin sheath* genes (GO:0043209, *p* = 42) and *astrocyte differentiation* genes (GO:0048708, *p* = 72), primarily functioning in excitatory neurons, oligodendrocytes and astrocytes, respectively. Specifically, the *excitatory synapse* gene set contains genes whose products function mainly in excitatory synapses, and the *myelin sheath* gene set has genes related to myelin sheath, which is supplied by oligodendrocytes to the central nervous system; the *astrocyte differentiation* gene set contains genes involved in the differentiation process of an astrocyte. These gene sets, according to their GO definitions, are expected to express and/or co-express primarily in the cell types that are relevant to their functions. In our analysis of these three gene sets, we focused on genes expressed in more than 25% of the ROSMAP bulk samples, resulting in sets of sizes *p* = 45, 41 and 68, respectively.

Figure 3 shows the co-expression estimates from `Bulk`, `CSNet` and `bMIND` for the *excitatory synapse* gene set. It is seen that `CSNet` identified co-expressions specific to excitatory neurons, while `bMIND` suggested similar co-expression patterns in all four cell types. We also estimated cell-type-specific co-expressions for the *myelin sheath* and *astrocyte differentiation* gene sets, shown in Figures S4 and S5, respectively. These plots show that `CSNet` identified co-expressions specific to oligodendrocyte and astrocyte, respectively, while `bMIND` again estimated similar co-expressions across four cell types. Finally, Figure 4 shows that the estimates based on single cell data are noisy and do not show any cell-type-specific co-expression patterns. Also in Figure 4, for all three gene sets, the strongest co-expressions are always observed in excitatory neurons, likely driven by the fact that it is the most abundant cell type (Mathys et al., 2019).

**Figure 3:**
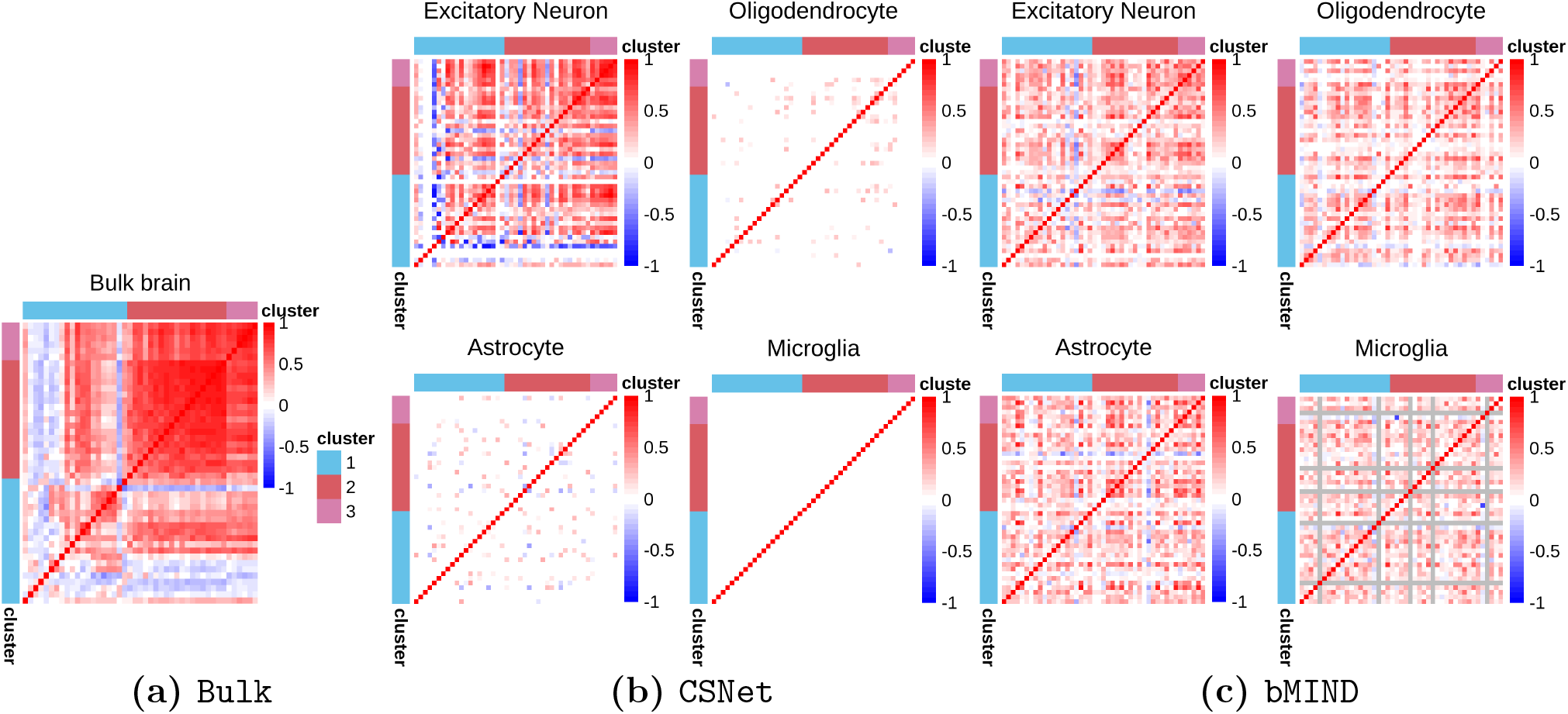
Co-expressions of *excitatory synapse* genes in different cell types. From left to right: (a) sample correlation matrix estimated from bulk RNA-seq data, (b) `CSNet` estimates and (c) `bMIND` estimates. For `bMIND`, genes with constant expression estimates across samples are marked in gray.

**Figure 4:**
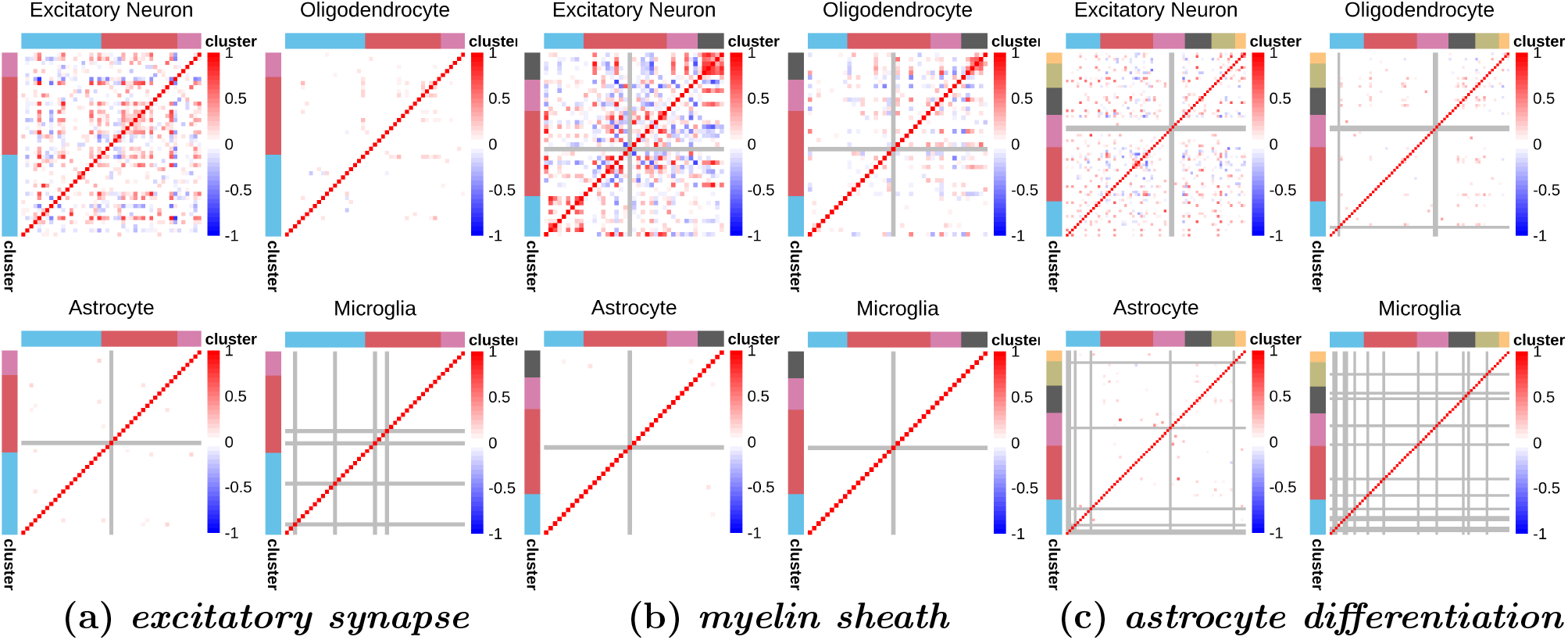
Co-expressions of *excitatory synapse, myelin sheath* and *astrocyte differentiation* genes in different cell types, estimated using the ROSMAP single cell data (Mathys et al., 2019). Marked in gray are genes with no variation or no expressions within a given cell type.

### 4.2 Alzheimer’s disease risk gene set

Next, we focused on Alzheimer’s disease risk genes from GWAS (see gene names in Table 3), which capture around 50% of the heritability in late-onset AD (Sims et al., 2020). Our analysis focused on 61 genes with a FPKM greater than 0.1 in at least 50 ROSMAP samples. There is a growing literature on the molecular mechanisms and related cell types for these risk genes. Besides the well studied pathways of amyloid-*β* and tau processing, several other pathways have also been implicated (Pimenova et al., 2018; Sims et al., 2020), among which neuroinflammation was recently highlighted as one of the most important causal pathways in Alzheimer’s disease (Heneka et al., 2015). Both microglia and astrocyte are the key cell types involved in such immune responses, and microglia, the innate immune cells in central nervous system, were prioritized as the cell type most enriched for GWAS associations (Skene and Grant, 2016; Tansey et al., 2018). Our analysis aims to use `CSNet` to explore the celltype-specific co-expression patterns among these Alzheimer’s disease risk genes.

**Table 3:** The list of Alzheimer’s disease risk genes. Gene names are displayed by cluster and in the same order as they appear in Figure 5. Our analysis included only the sequenced genes with an FPKM greater than 0.1 in at least 50 ROSMAP samples.

| Cluster | Genes |
| --- | --- |
| 1 | MEF2C, PILRA, MS4A8, EPHA1, IL34, CELF1, KAT8, KANSL1, HS3ST1, ACE, PTK2B, MAPT, NDUFAF7, TSPOAP1-AS1, SLC24A4, SORL1, PPARGC1A, C17orf107, APP, TPBG, CNTNAP2 |
| 2 | IGHG3, HLA-DRB1, TRIP4, CASS4, MS4A14, PLCG2, INPP5D, SCIMP, MS4A7, CD33, TREM2, MS4A4A, MS4A6A, ABI3, SPI1 |
| 3 | PRKD3, TP53INP1, CD2AP, ZNF423, CLU, SPPL2A, FERMT2, IQCK, RORA, ECHDC3 |
| 4 | APOE, MS4A4E, ZCWPW1, ABCA7, BIN1, SHARPIN |
| 5 | HESX1, APH1B, ADAM10, PICALM, ZNF655 |
| 6 | CR1, HBEGF, ADAMTS1, ADAMTS4 |

Figure 5 shows the estimates from `Bulk`, `d-CSNet` and `CSNet`, respectively. The `bMIND` estimates are again similar across cell types, and are relegated to Figure S6(a) in the supplement. In Figure 5, some within cluster co-expressions from bulk samples are no longer seen in the cell-type-specific estimates, likely due to the confounding effect of cell type proportions. The `CSNet` estimates in Figure 5(c) show that Cluster 4 (colored in black) were highly coexpressed in astrocytes. This gene cluster includes APOE, a major Alzheimer’s disease risk gene known to be highly expressed in astrocytes (Yamazaki et al., 2019). APOE protein is primarily produced in astrocytes, which then interacts with amyloid-*β*, which is involved in a central pathway of Alzheimer’s disease (Yamazaki et al., 2019). Besides, both APOE and ABCA7 contribute to lipid metabolism and phagocytosis (Pimenova et al., 2018), consistent with their high co-expressions found in Cluster 4. The `CSNet` estimates for Cluster 4 further highlight their connections with several other Alzheimer’s disease risk genes in astrocytes. Additionally, the `d-CSNet` estimates in Figure 5(b) suggest Cluster 2 (colored in red) were co-expressed in microglia, though the signals are relatively weak and the `CSNet` estimator for microglia is nearly diagonal. Nevertheless, the co-expression in Cluster 2 is likely microglia specific supported by several existing findings in Alzheimer’s disease. Firstly, 9 out of 15 genes in Cluster 2 are known to be involved in neuroinflammation and Alzheimer’s disease pathology via microglia. Among them, the coding variants in PLCG2, TREM2, ABI3 implicate innate immunity in Alzheimer’s disease as mediated by microglia (Sims et al., 2017); CD33 inhibits the uptake of amyloid-*β* in microglia (Griciuc et al., 2013); MS4A gene cluster is a key modulator of TREM2 in microglia (Deming et al., 2019) and SPI1 is a central regulator of microglia expression and Alzheimer’s disease risk (Kosoy et al., 2021). In addition, 9 genes are known to express uniquely in microglia, including HLA-DRB1, PLCG2, CD33, TREM2, ABI3 and the MS4A gene cluster (Sims et al., 2017; Pimenova et al., 2018). The `d-CSNet` estimates were able to identify cell-type-specific co-expression patterns of these genes, while single cell data based estimates could not [see Figure S6(b)], and possibly offer new insights into regulations of Alzheimer’s disease risk genes. The estimated co-expressions in gene Clusters 1 and 2 reveal previously unknown cell-type-specific co-expressions among Alzheimer’s disease risk genes, and may suggest cell-type-specific disease pathology for AD.

**Figure 5:**
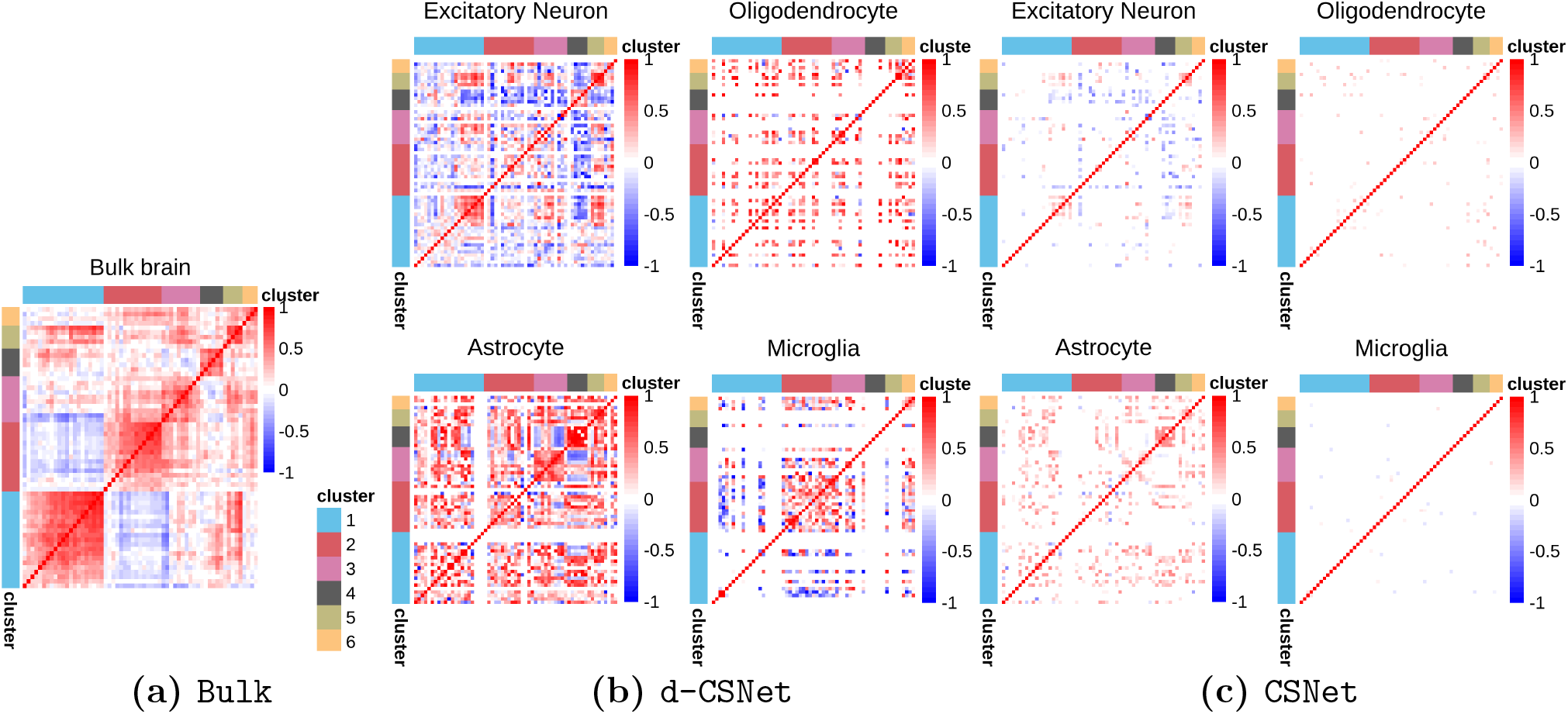
Co-expression networks of Alzheimer’s disease risk genes inferred from the ROSMAP data. From left to right: sample correlation matrix estimated from bulk RNA-seq data [panel (a)], d-CSNet estimates [panel (b)], and CSNet estimates [panel (c)].

Finally, the sensitivity analysis in Section A2.1 shows that `CSNet` remains robust as a reasonable amount of noise is added to the cell type proportions. We have also added a negative control experiment where cell type proportion vectors for different samples were randomly permuted. Figure S8 shows that the resulting estimates in excitatory neurons, the most abundent cell type, always resemble the bulk co-expression estimate, while the previously uncovered cell-type-specific co-expression patterns are no longer seen.

## 5 Theoretical Properties

In this section, we establish the non-asymptotic convergence rate of `CSNet`, the sparse celltype-specific correlation estimates, and also establish variable selection consistency, ensuring that we correctly identify edges in the cell-type-specific co-expression networks with probability tending to 1. Our theoretical analysis is challenging with a few unique aspects. First, we assume the expression profile from each cell type follows (marginally) a sub-exponential distribution to accommodate the commonly used negative binomial distributions in modeling read counts from RNA-seq data (Robinson et al., 2010). In this case, each element *z*_*ij*_*z*_*ij*′_ in the response vector in (8) is the product of two sub-exponential random variables, which requires a new concentration result (see Lemma A3.2 and its proof in Section A3.4). Second, our procedure considers correlation estimates, which are normalized with estimated variances. Thus, a more delicate analysis is needed to find the non-asymptotic convergence rate of 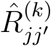 to 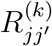. Third, as ***μ***^(*k*)^’s are unknown, the response 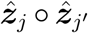 in (8) inherits errors from estimating the mean parameter ***μ***^(*k*)^’s. Next, we state a few regularity conditions.

### Assumption 1.

*Let* 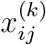 *follow a sub-exponential distribution, i* ∈ [*n*], *j* ∈ [*p*], *k* ∈ [*K*] *and* 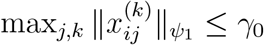 *for some positive constant γ*_0_, *where* 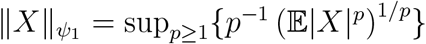.

Assumption 1 imposes a marginal distribution assumption on 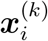, the expression profile from the *k*th cell type in sample *i*, in that each element 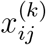 follows a sub-exponential distribution with a bounded sub-exponential norm. The sub-exponential assumption is more relaxed than the Gaussian or sub-Gaussian assumption commonly imposed (Bickel and Levina, 2008a,b; Rothman et al., 2008, 2009) and includes distributions such as negative binomial, which is often used to model read counts from RNA-seq data (Robinson et al., 2010).

### Assumption 2.

*Let* ***R***^(*k*)^ ∈ *𝒰* (*q, s*_*p*_) *for* 0 *≤ q<* 1, *where 𝒰* (*q, s*_*p*_) *is a class of approximately sparse matrices with sparsity parameter s*_*p*_ *(can be a function of p), defined as*

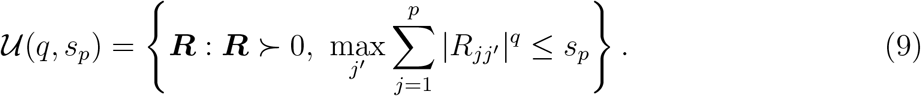

This assumption stipulates that the cell-type-specific correlation matrices are approximately sparse. The definition of approximate sparsity in (9) is commonly employed in large covariance matrix estimation (e.g., Bickel and Levina, 2008a; Rothman et al., 2008). When *q* = 0, Assumption 2 imposes that the number of non-zero entries in each column of ***R***^(*k*)^ is less than *s*_*p*_, *k* ∈ [*K*]. In Assumption 2, the sparse parameters *q* and *s*_*p*_ may vary across *k*, i.e., 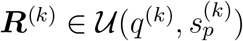, though we dropped the superscript *k* to simplify notation, which can be viewed as taking *q* = max_*k*_ *q*^(*k*)^ and 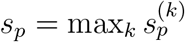.

Define two matrices **Ψ**_*K×K*_ and **Φ**_*K×K*_, where 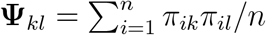 and 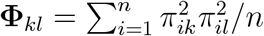, for *k, l* ∈ [*K*]. It then holds that **Ψ**_*K×K*_ = ***D***^T^***D****/n* and **Φ**_*K×K*_ = ***H***^T^***H****/n*, where ***D*** and ***H*** are as defined in (5) and (6), respectively.

### Assumption 3.

*There exist constants* 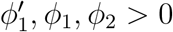 *such that the eigenvalues of **Ψ***_*K×K*_ *are lower-bounded by* 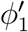, *the eigenvalues of* **Φ**_*K×K*_ *are lower-bounded by cp*_1_ *and* min_*k*_ **Φ**_*kk*_ ≥ *ϕ*_2_.

Assumption 3 places two regularity conditions on the cell type proportion matrices ***D, H*** ∈ ℝ^*n×K*^, which can be viewed as the fixed design matrices in (3) and (4). The bounded eigenvalue conditions on **Ψ**_*K×K*_ and **Φ**_*K×K*_ are not restrictive as the number of cell types *K* is treated as fixed. The condition min_*k*_**Φ**_*kk*_ *≥ ϕ*_2_ stipulates that the average proportion for each cell type (over all bulk samples) should be bounded away from zero, which is a mild condition. In fact, if the cell type proportions are from an underlying multinomial or Dirichlet distribution with fixed parameters, all conditions in Assumption 3 are expected to hold; see proof in Section A3.6.

### Theorem 5.1.

*Let* 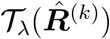 *be the estimator as defined in* (8). *Under Assumptions 1-3*, log *p* = *o*(*n*^1/3^) *and* 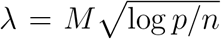 *for a sufficiently large constant M* > 0, *we have, for k* ∈ [*K*],

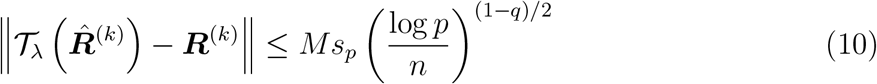

*with probability at least* 1 − *C*_1_ exp(− *C*_2_ log *p*), *where C*_1_ *and C*_2_ *are some positive constants*.

Theorem 5.1 implies that the number of genes *p* can far exceed the sample size *n*, as long as *s*_*p*_(log *p/n*)^(1−*q*)/2^ tends to zero. The condition log *p* = *o*(*n*^1/3^) is needed as 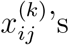 are assumed to follow a sub-exponential distribution. When *q* = 0, the convergence rate in (10) further simplifies to 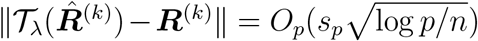, which is comparable with the convergence rate derived in estimating sparse covariance matrices (e.g., Rothman et al., 2008). Based on (10) and using a similar argument as in Bickel and Levina (2008a), it can be shown that the convergence rate in Frobenius norm is 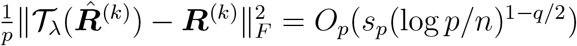.

Denote the support of a matrix ***R*** by Ω(***R***) = {(*j, j*′) : *R*_*jj*′_ *≠* 0}. We show that the support of ***R***^(*k*)^ is recovered with high probability assuming a minimal signal condition.

### Theorem 5.2.

*Suppose Assumptions 1 and 3 hold. Let* 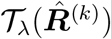 *be the estimator as defined in* (8), 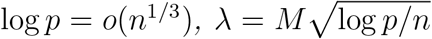 *for a sufficiently large constant M >* 0. *If* 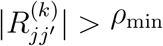 *for some ρ*_*min*_ *>* 0 *such that* 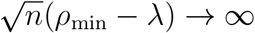, *then we have for k* ∈ [*K*]

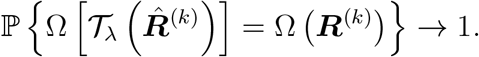

Theorem 5.2 holds without the approximately sparse condition in Assumption 3, though it requires a minimal signal condition on the nonzero elements in ***R***^(*k*)^’s. Such a condition is commonly considered in establishing selection consistency (e.g., El Karoui, 2008) and our result allows the minimal signal to tend to zero as *n* increases (e.g., *ρ*_min_ = 2 *λ*). Theorem 5.2 also has an important implication in practice as it ensures that the selected set of edges after thresholding is the same as the true set of edges with probability tending to one.

## 6 Discussion

We conclude the article with a few remarks. First, the model (1) is designed for gene expression data measured by the RNA-seq protocol, where sequencing read counts capture the expression levels for all cells in a tissue sample. We caution that the same model may not be applicable to microarray data, where expression levels have been transformed for normalization (Zhong and Liu, 2012). In (1), 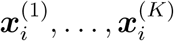 are assumed independent, as we focus on cell types with distinct functions and gene expression profiles across different cell types are likely not highly correlated (see Figure S7.) When this assumption does not hold, we can expand the covariance expression in (2) to include the cross product terms Σ_(*k*≠*k*′)_ *π*_*ik*_*π*_*ik*′_Σ^(*k,k*′)^, where Σ^(*k,k*′)^ ∈ ℝ^*p×p*^ is the cross-covariance 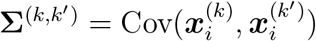. In this case, the estimation procedure can be carried out similarly, though the total number of parameters increase from *O*(*Kp*^2^) to *O*(*K*^2^*p*^2^).

Next, we have assumed that the cell-type proportions *π*_*ik*_’s are given in our analysis. For example, they can be inferred using existing methods such as CIBERSORTx (Newman et al., 2019). Our empirical investigations showed that `CSNet` is not overly sensitive to errors in *π*_*ik*_’s (see sensitive analysis in Section 3.2 and A2.1). It is possible to further extend our framework to accommodate noisy *π*_*ik*_’s. In this case, we may further consider the 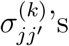 to be estimated from 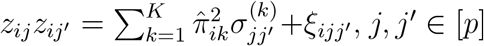, where 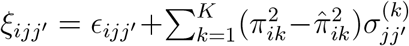. Hence, if the error 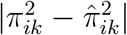 is small, 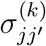 should still be well estimated. We leave the full investigation of this topic as future research. As the correlation matrices are estimated by element and then thresholded, the correlation matrix estimates are not guaranteed to be positive definite. However, as the true correlation matrices are positive definite, it follows from Theorems 5.1-5.2 that the estimated correlation matrices are asymptotically positive definite. For finite sample cases, it may be desirable to ensure the positive definiteness of the final estimator. One strategy is to solve a constrained optimization problem, subject to positive definiteness, to find the nearest correlation matrix in Frobenius norm. This can be carried out efficiently using existing solvers (e.g., Higham, 2002).

Finally, we have focused on using bulk samples for co-expression analysis. With an increasing number of studies collecting single cell data, we may obtain more accurate coexpression estimates through integrated analysis of bulk and single cell data. For example, Morabito et al. (2021) and bMIND have explored ways to extract information from single cell data to help with the estimation. However, platform differences and batch effects are prominent in integration, and have not been addressed well in these methods. We plan to explore along this direction in our future research.

## Acknowledgements

We thank the ROSMAP project for their permission, requested at https://www.radc.rush.edu, to access the bulk RNA-seq and single nueclues RNA-seq data in the project. The ROSMAP project is supported by the following grants: P30AG72975, P30AG010161 (ADCC), R01AG015819 (RISK), R01AG017917 (MAP), U01AG46152 (AMP-AD Pipeline I) and U01AG61356 (AMP-AD Pipeline II).

## Supplementary Materials

### A1 Simulating from a multivariate negative binomial distribution with a large and structured covariance

Consider a negative binomial (NB) random variable distributed as

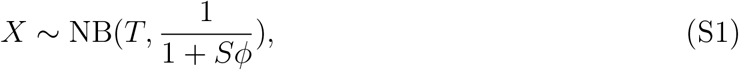

such that

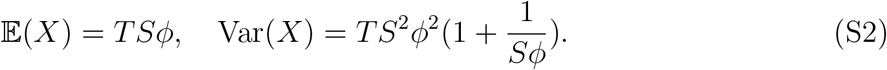

Our procedure simulates a Poisson-Gamma mixture that follows (S1) by simulating

- Gamma: *θ* ∼ Gamma(*T, ϕ*), where 𝔼[*θ*] = *Tϕ* and Var[*θ*] = *Tϕ*^2^;
- Poisson: *X*|*θ, S* ∼ Poisson(*Sθ*).

This is a model commonly employed in differential expression analysis methods for bulk RNA-seq data (Robinson et al., 2010; Love et al., 2014), where *θ* models gene abundance, *S* denotes the total number of mRNA transcripts (i.e., sequencing depth) and *X*|*θ* models the procedure of sampling a transcript from the pool. The interpretation of parameter *T* will be explained in the following paragraph. In addition, we note that this model (S1) sets the variance to be a quadratic function of the mean, which is supported by observations on real bulk RNA-seq data (Chen et al., 2014).

Before we proceed, we first discuss a useful property of Gamma random variables which entails that if *g*_1_ and *g*_2_ are two independent samples from Gamma(*T, ϕ*), then *g*_1_ + *g*_2_ ∼ Gamma(2*T, ϕ*). This property facilitates an effective procedure for generating correlated Gamma random variables. For example, if *g*_1_, *g*_2_, and *g*_3_ are independent Gamma(*T, ϕ*) random variables, then Cor(*g*_1_ + *g*_2_, *g*_2_ + *g*_3_)=0.5, with *g*_2_ being the component that is being shared by two Gamma random variables *g*_1_+*g*_2_ and *g*_2_+*g*_3_. This idea of simulating correlated Gamma random variables has previously been explored by Ronning (1977).

#### Algorithm 1 Algorithm for simulating multivariate negative binomial random variables

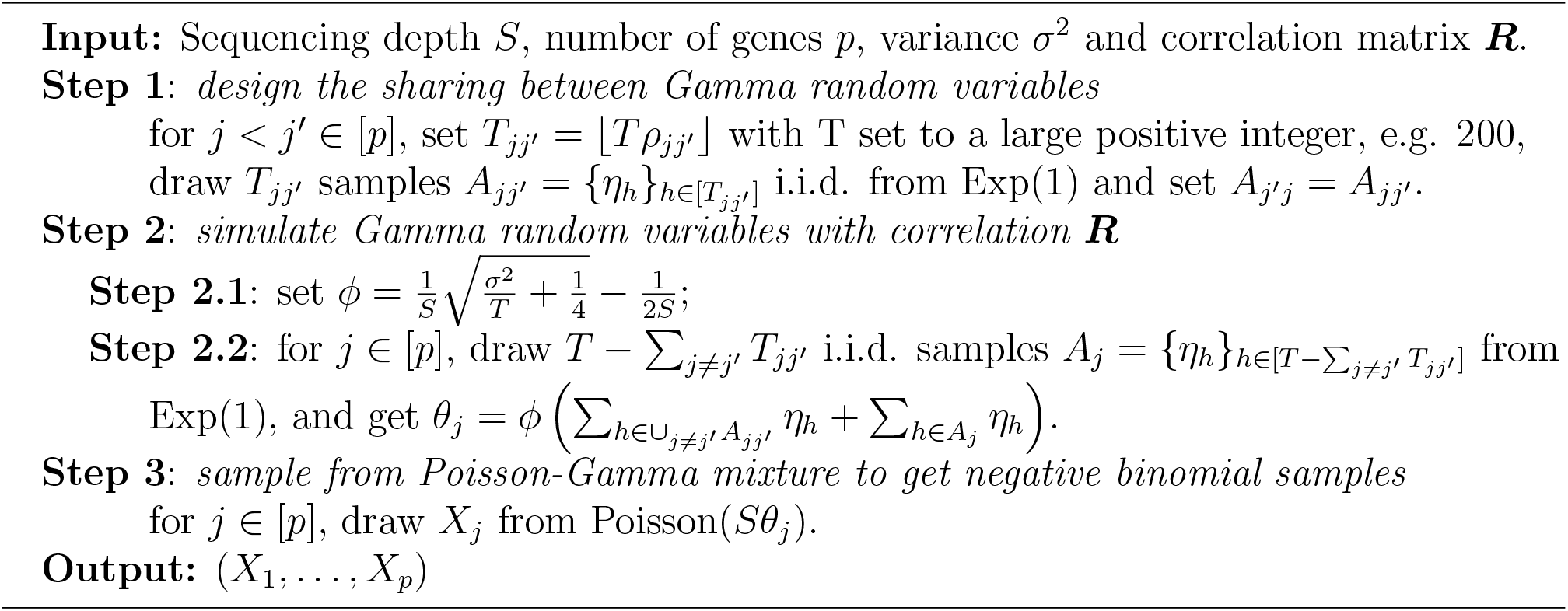

Our proposed procedure that simulates correlated negative binomial random variables, detailed in Algorithm 1, is divided into the following three steps. Step 1 designs the sharing between Gamma random variables, Step 2 simulates correlated Gamma random variables and Step 3 samples from the Poisson-Gamma mixture to get desired negative binomial samples. Specifically, in Algorithm 1, Step 1 sets *T* to be a large constant, whose subsets will be shared across samples as specified in Step 2.2, Step 2 draws ‘building blocks’ of Exp(1) random variables (i.e., Gamma(1,1)) to be shared and then generates correlated Gamma random variables and Step 3 employs Poisson sampling to obtain negative binomial random variables. We note that the assumption of homogeneous variances across all genes can be relaxed by allowing *ϕ* to vary across samples.

The output of Algorithm 1 is a *p*-dimensional random vector (*X*_1_, …, *X*_*p*_) with Var[*X*_*j*_] = *σ*^2^ for *j* ∈ [*p*] and correlation matrix ***R*** *×* [1*/*(1 + 1*/Sϕ*)]. It is seen that the correlation is biased from the specified correlation matrix ***R*** by a multiplicative factor *b* = 1*/*(1 + 1*/Sϕ*). This factor is close to 1 when 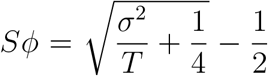 is large, as implied by (S2). Therefore, this bias is negligible for genes with large variances. Our own analysis of real bulk RNA-seq data found *S* = 6 *×* 10^7^ to be a common sequencing depth, and *σ* ^2^ = exp(8) to be representative of gene variances for highly-expressed protein coding genes. Under this setting, the multiplicative bias *b* = 0.77. Correspondingly, in Section 3 we simulate MA(1) structure with *ρ* = 0.5 *× b ≈* 0.39 and an AR(1)-type structure with *ρ*_*jj* ′_ = 0.70 *×* 0.9^|*j − j* ′ *−* 1|^ for *j ≠ j*′, where 0.70 *≈* 0.9 *× b*. Finally, as an example, Figure S9 demonstrates that Algorithm 1 simulates samples that are faithful to the prescribed AR(1)-type correlation structures in Figure 1.

### A2 More Results from Data Analysis in Section 4

#### A2.1 Sensitivity analysis

For the four gene sets in Section 4, we conduct a sensitivity analysis to evaluate the robustness of the `CSNet` estimates against perturbations in the cell type proportions inferred using CIBERSORTx. For each sample *i* and given ***π***_*i*_, the inferred cell type proportions using CIBERSORTx, the perturbed cell type proportions 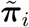 are independently simulated from Dirichlet(***π***_*i*_ *× k*), where *k* is set to {10, 100, 1000}. A smaller *k* corresponds to a larger noise level, as illustrated in Figure S1. Furthermore, we evaluate the difference between the 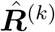, `CSNet` estimated with ***π***_*i*_, and 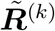, `CSNet` estimated with 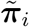, by Frobenius norm measured as 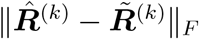, TPR measured as 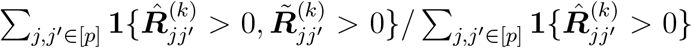 and FPR measured as 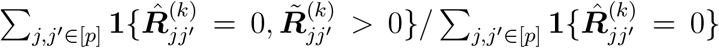. Figure S2 shows that `CSNet` estimates are robust under a reasonable amount of noise.

**Figure S1:**
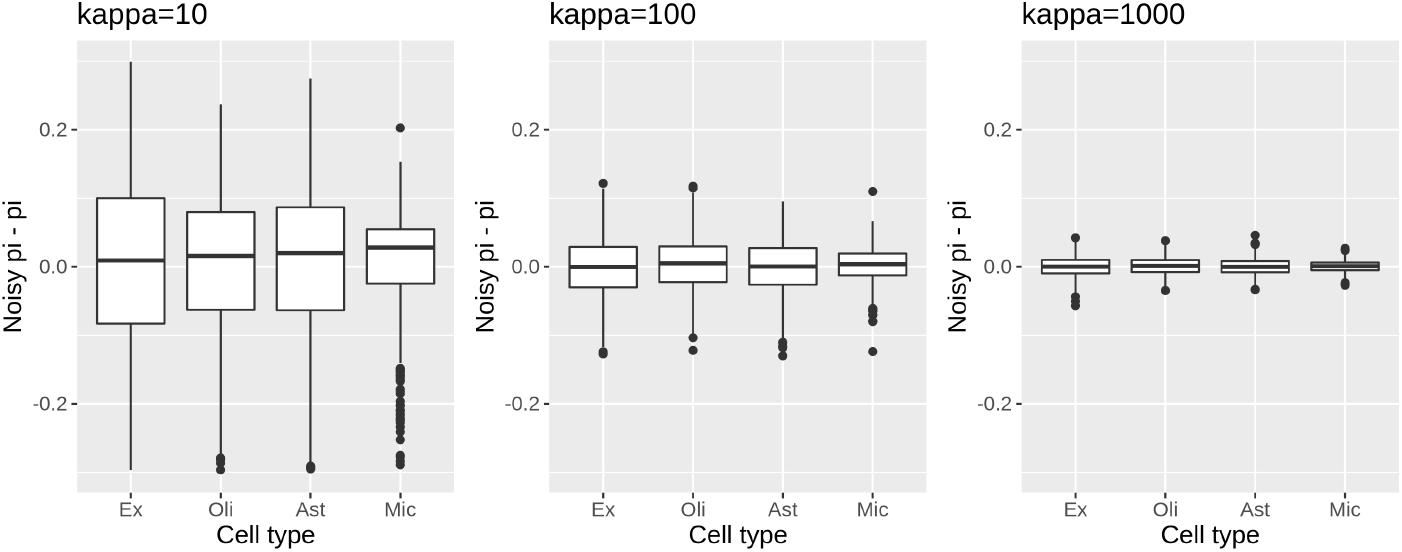
Boxplots of 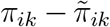 for four most abundant cell types: excitatory neuron (Ex), oligodendrocyte (Oli), astrocyte (Ast), microglia (Mic).

**Figure S2:**
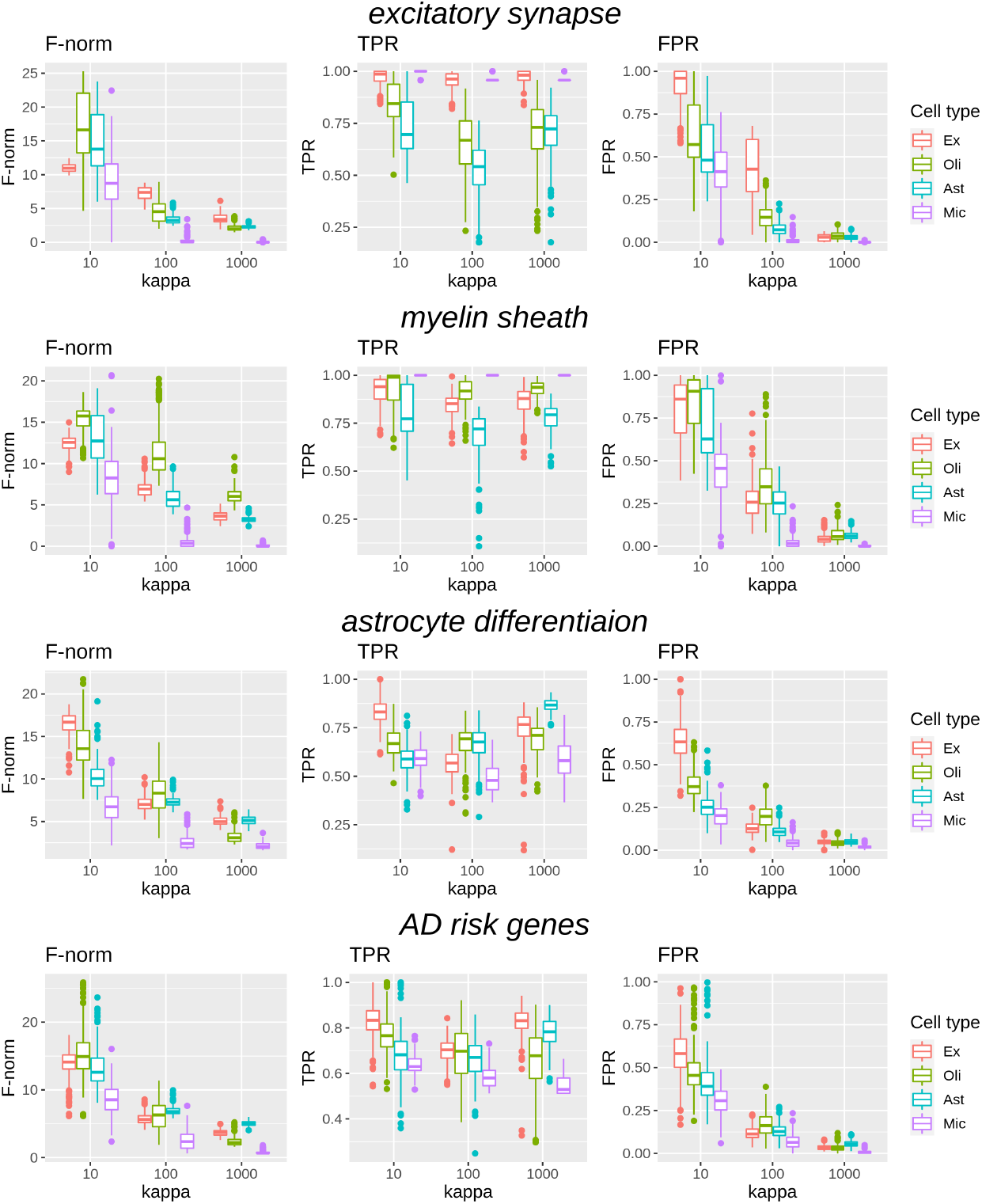
Evaluation criteria with varying noise level *k* for four gene sets. From top to bottom: *excitatory synapse, myelin sheath, astrocyte differentiation* and Alzheimer’s disease (AD) risk genes. Results are shown for four most abundant cell types: excitatory neuron (Ex), oligodendrocyte (Oli), astrocyte (Ast), microglia (Mic).

#### A2.2 Additional figures

**Figure S3:**
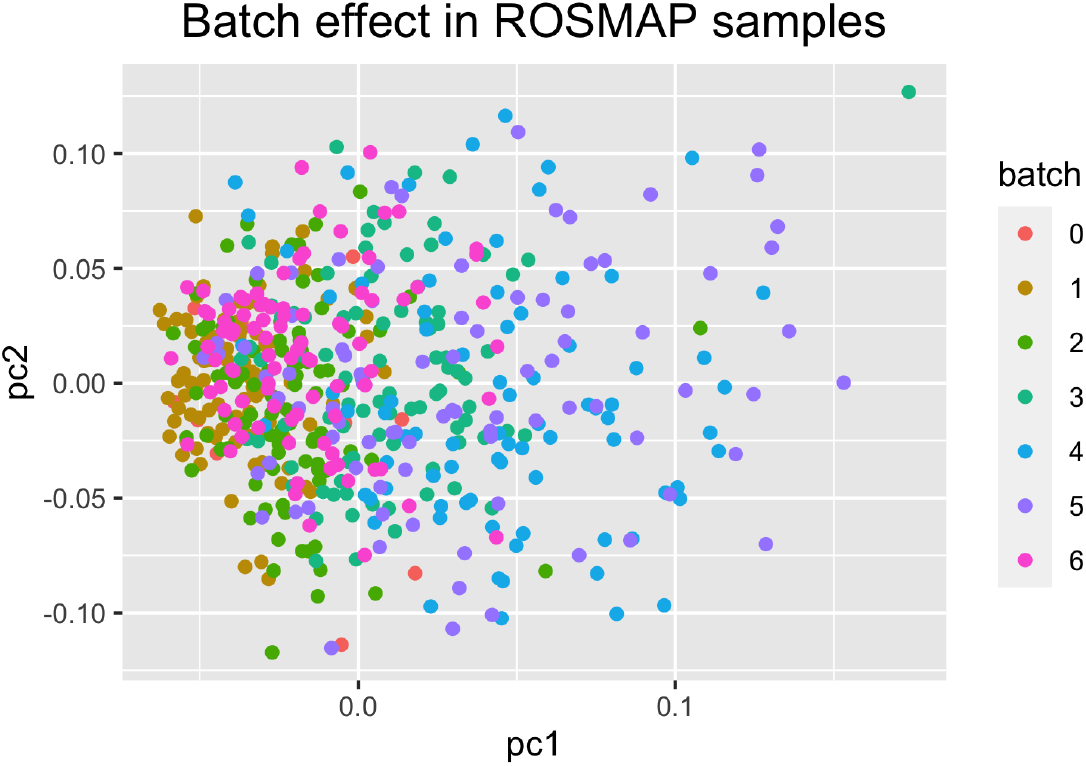
Top two PCs from the principal component analysis of the 541 ROSMAP samples using protein coding genes with average FPKM*>* 0.25, with points colored by processing batches.

**Figure S4:**
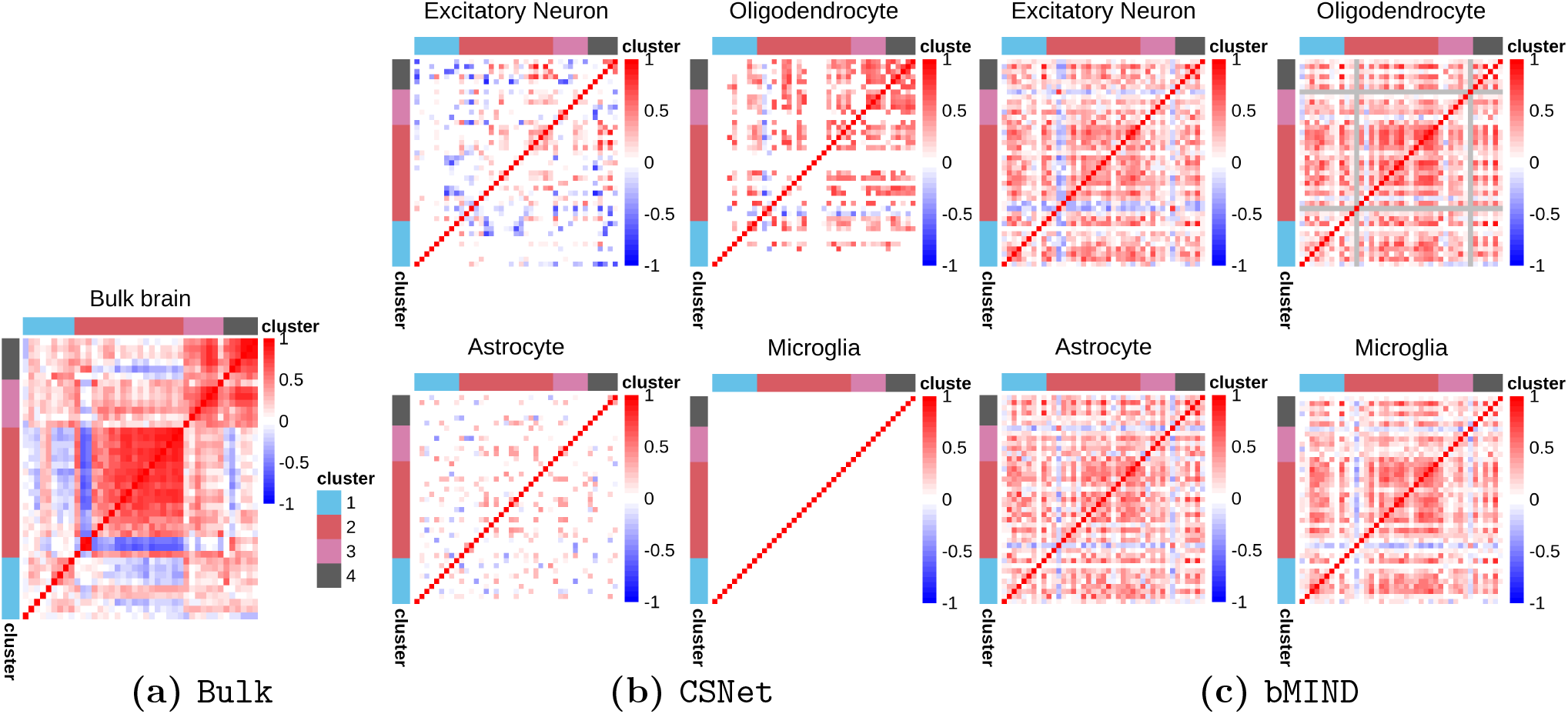
Co-expressions of *myelin sheath* genes in different cell types. From left to right: (a) sample correlation matrix estimated from bulk RNA-seq data, (b) `CSNet` estimates and (c) `bMIND` estimates. For `bMIND`, genes with constant expression estimates across samples are marked in gray.

**Figure S5:**
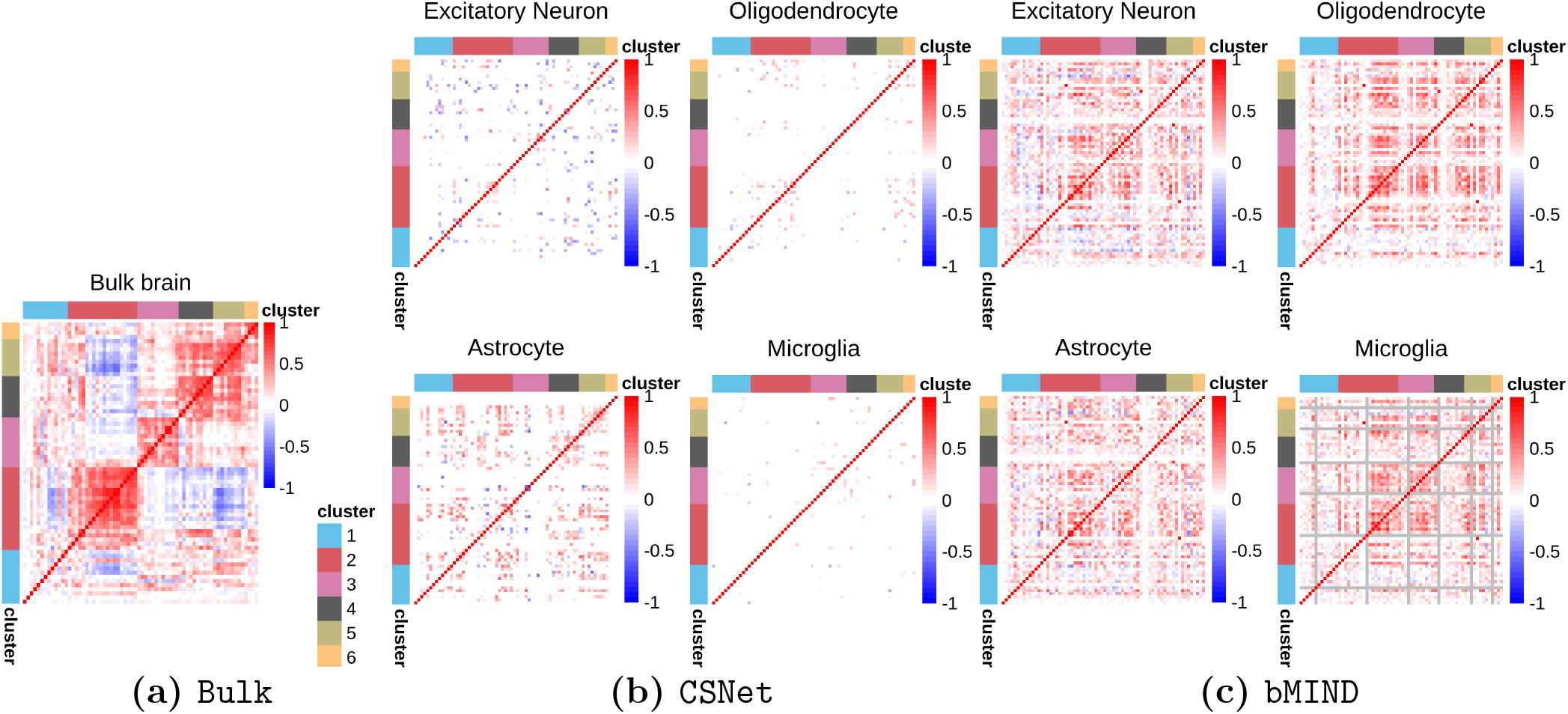
Co-expressions of *astrocyte differentiation* genes in different cell types. From left to right: (a) sample correlation matrix estimated from bulk RNA-seq data, (b) `CSNet` estimates and (c) `bMIND` estimates. For `bMIND`, genes with constant expression estimates across samples are marked in gray.

**Figure S6:**
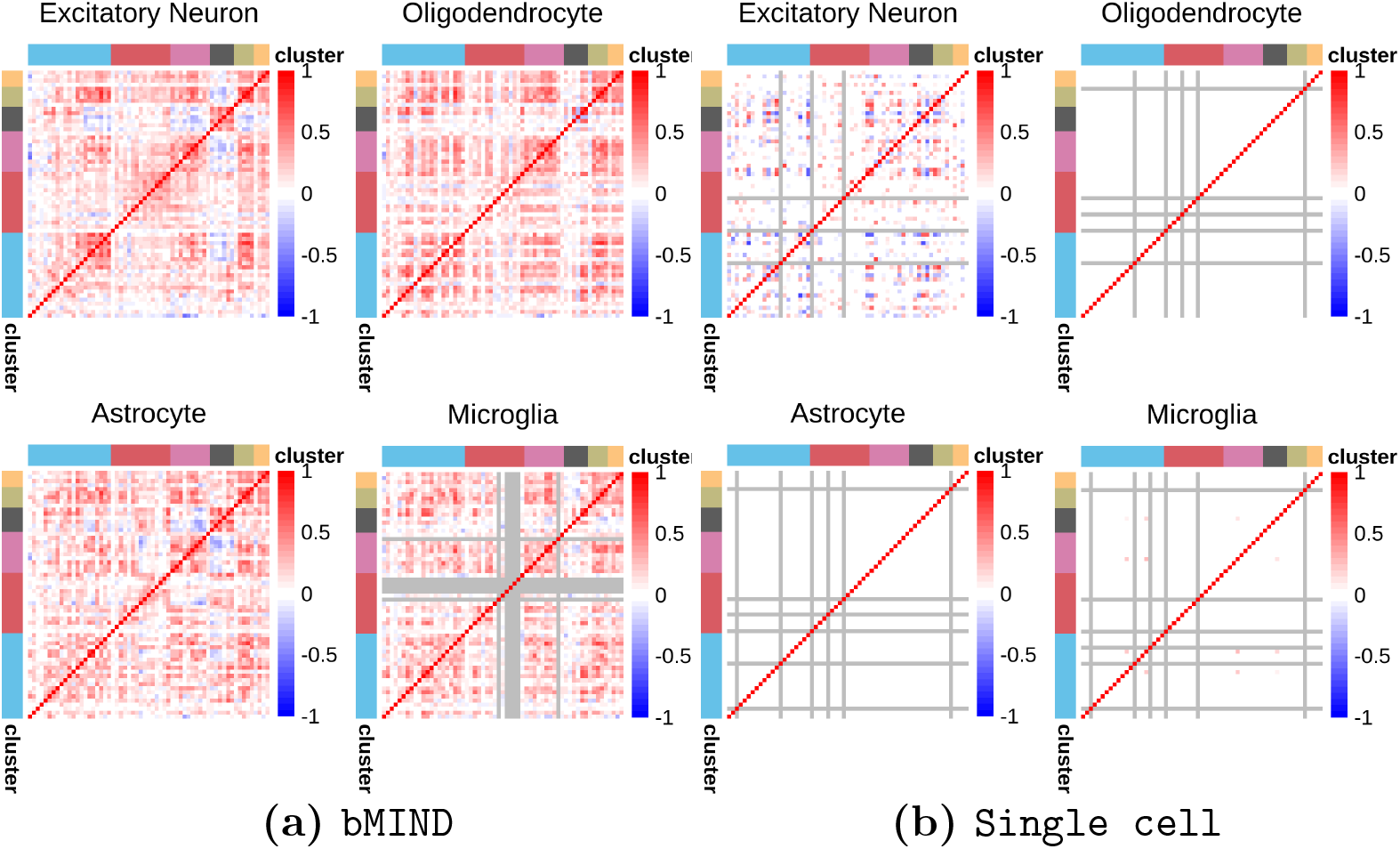
Co-expressions networks of Alzheimer’s disease risk genes estimated with (a) `bMIND`, (b) single cell data from Mathys et al. (2019). Genes with constant expression estimates across samples are marked in gray.

**Figure S7:**
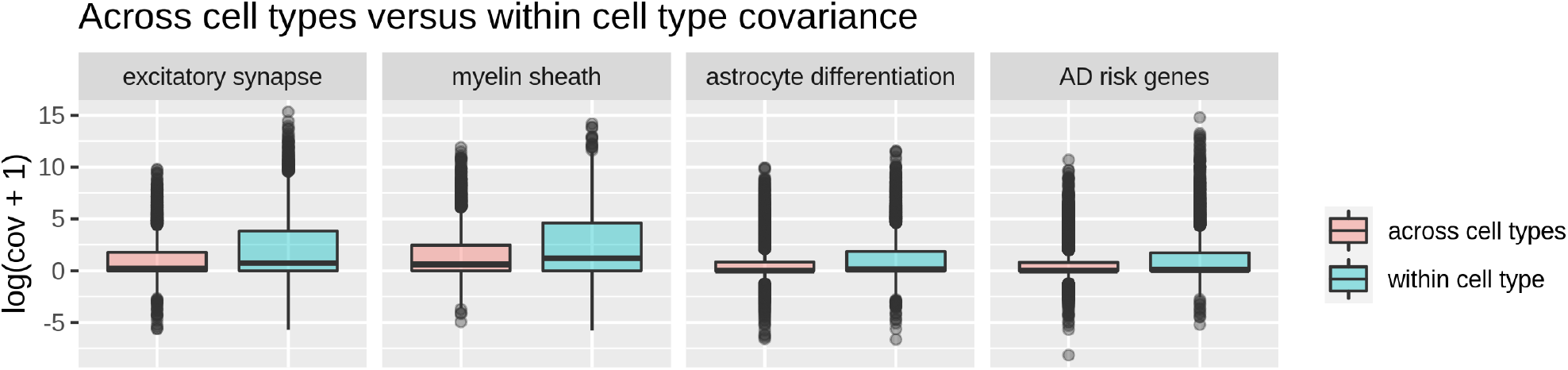
Cross-cell-type covariances Cov 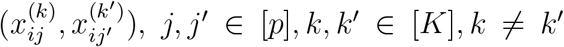 and within-cell-type covariances Cov 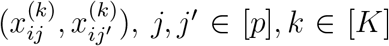 for four gene sets, evaluated with single cell data from Mathys et al. (2019) following a similar procedure as described in Section 4.

**Figure S8:**
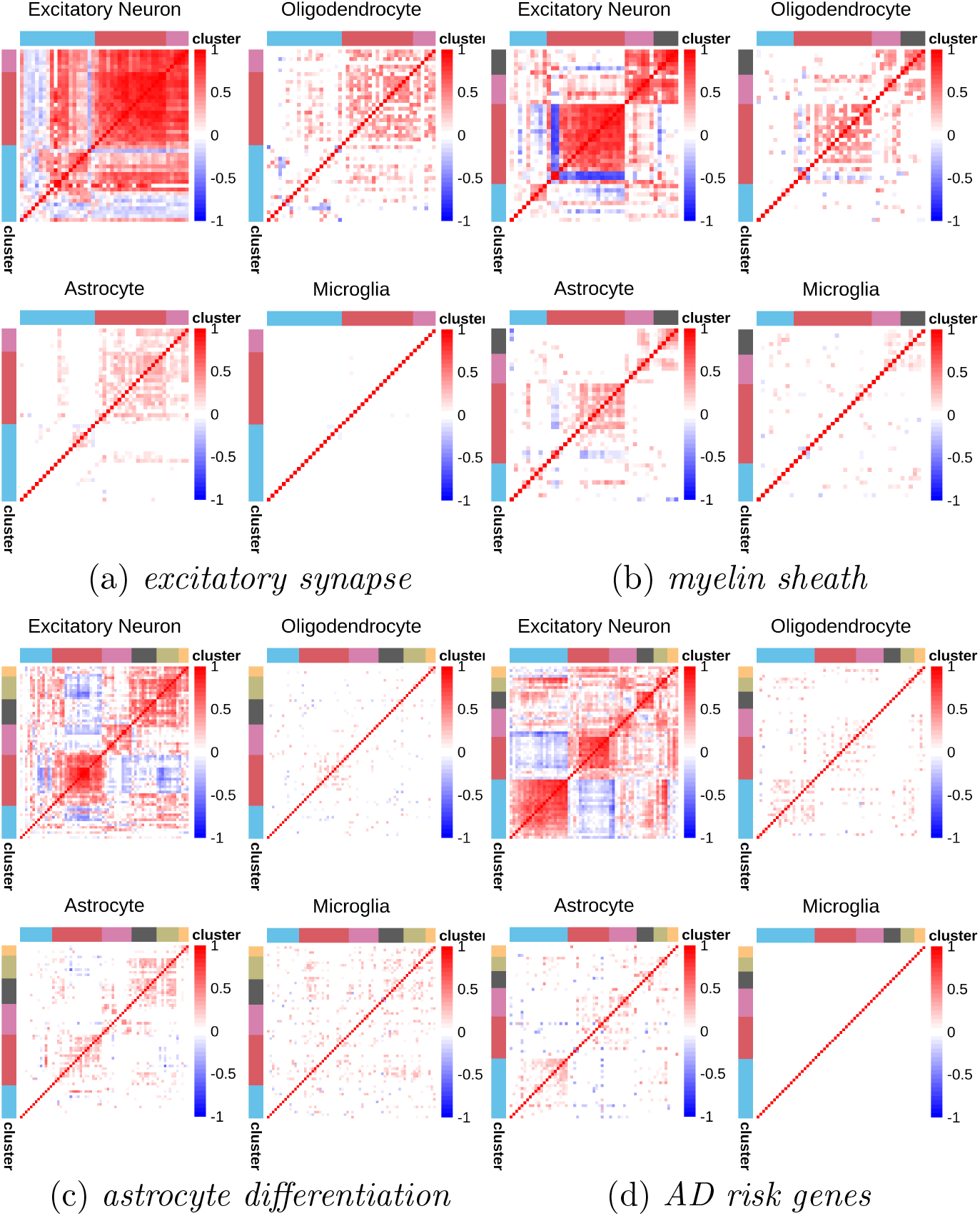
`CSNet` estimates of co-expressions in the *excitatory synapse, myelin sheath, astrocyte differentiation* and Alzheimer’s disease (AD) risk genes gene sets, with proportions randomly permuted across samples.

### A3 Proofs of Main Results

#### A3.1 Technical Lemmas

We first present a set of technical lemmas. The proofs for Lemmas A3.2 and A3.5 are presented in Sections A3.4 and A3.5, respectively.

##### Lemma A3.1.

*[Theorem 2*.*8*.*11 in Vershynin (2018)] Let* {*X*_1_, …, *X*_*n*_} *be independent mean zero sub-exponential random variables with* 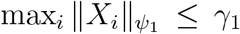 *for some constant γ*_1_ *>* 0. *Given* ***a*** = (*a*_1_, *···, a*_*n*_) ∈ ℝ^*n*^, *for every t>* 0, *we have*

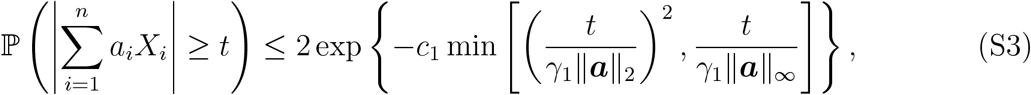

*for some positive constant c*_1_.

##### Lemma A3.2.

*Let* {*X*_1_, …, *X*_*n*_} *and* {*Y*_1_, …, *Y*_*n*_} *be two sets of independent sub-exponential random variables with* 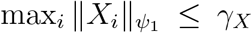 *and* 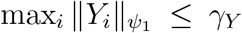, *respectively. Let* ***a*** = (*a*_1_, …, *a*_*n*_) ∈ ℝ^*n*^ *and γ*_2_ = γ_*X*_ *γ*_*Y*_. *Then there exist some constants d*_1_, *d*_2_, *c*_2_ *>* 0 *such that for* 0 *<t < d*_1_ *× γ*_2_ *× ‖****a****‖* _*∞*_,

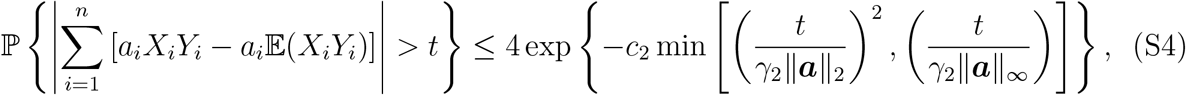

*and for any t ≥ d*_2_ *× γ*_2_ *× ‖****a****‖*_*∞*_,

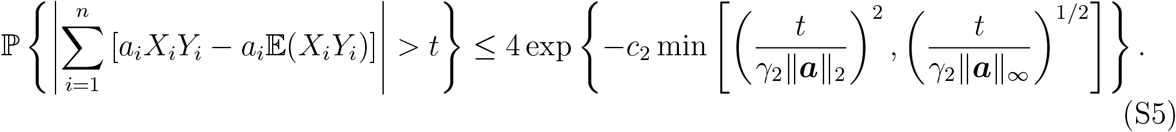

##### Lemma A3.3.

*[Theorem 1*.*4 Adamczak and Wolff (2015)] Let X* = (*X*_1_, …, *X*_*n*_) *be a random vector with independent components, such that for all* 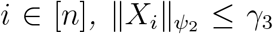, *where* 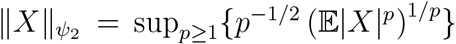. *Let 𝒫*_*d*_ *denote the set of partitions of* {1, …, *d*} *into nonempty, pairwise disjoint sets and ‖·‖* _*𝒥*_ *denote a tensor norm whose definition is relegated to Section A3.4. Then for every polynomial f:* ℝ^*n*^ *→* ℝ *of degree D and any t>* 0,

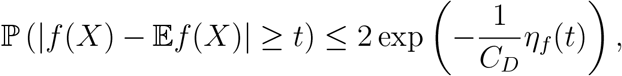

*where* 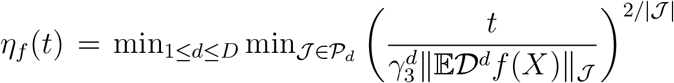, *𝒟*^*d*^*f denotes the d-th derivative of f and C*_*D*_ *is some positive constant depending on D*.

##### Lemma A3.4.

*[Lemma 1 and Theorem 1 Rothman et al. (2009)] Consider the class of sparse covariance matrices 𝒰* (*q, s*_*p*_) *as defined in Assumption 2. If* ***R*** ∈ *𝒰* (*q, s*_*p*_), *then the thresholding operator 𝒯 in* (8) *with threshold λ satisfies*

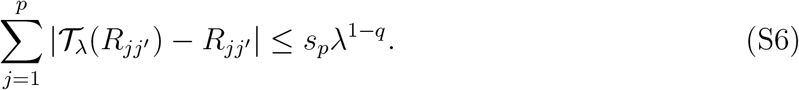

*In addition, suppose we have an estimator* 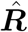 *such that*

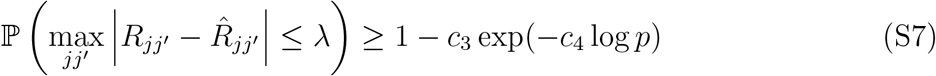

*for some λ >* 0 *and positive constants c*_3_ *and c*_4_. *Then*, 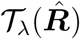 *satisfies*

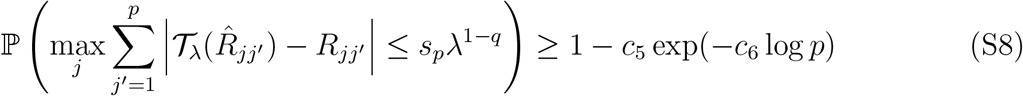

*for some positive constants c*_5_ *and c*_6_.

##### Lemma A3.5.

*Let X* = (*X*_1_, …, *X*_*n*_) *be a vector of sub-exponential random variables with* 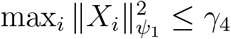. *Let* 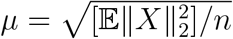. *Then, for t ≤ μ*,

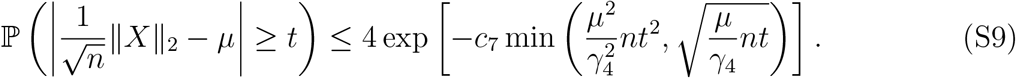

#### A3.2 Proof of Theorem 5.1

The proof is divided into three steps. In step 1, we establish an element-wise concentration inequality for the covariance estimates 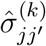. In step 2, we get an element-wise concentration inequality for the correlation estimates 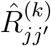. Step 3 puts together the previous steps and finds the appropriate thresholds that lead to the desired result.

**Step 1**. We first show that the element-wise covariance estimate 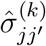 satisfies

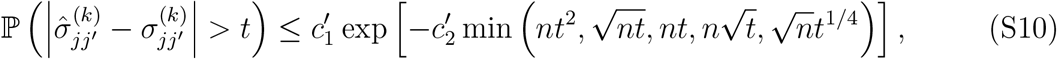

for *k* ∈ [*K*], *j, j*′ ∈ [*p*] and some positive constants 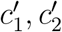.

Denote 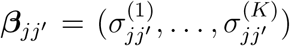. Based on (4) and (5), we have 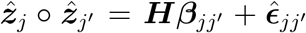, where 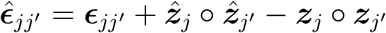, and ***z***_*j*_’s, 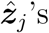 are as defined in Section 2. We consider the least squares estimates of covariances in (6), i.e. 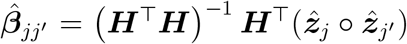, which satisfies

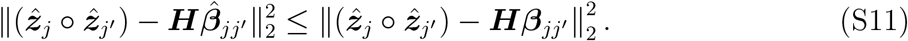

Recall that ***ϵ***_*jj*_′ = ***z***_*j*_ *°* ***z***_*j*′_ *−****Hβ***_*jj*′_. Letting 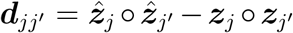 and 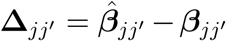, we have 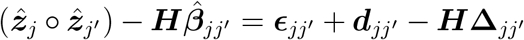 and 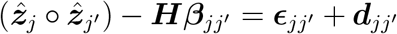. Then, (S11) gives that

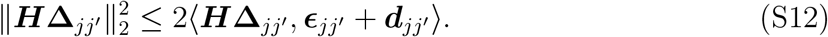

By Hölder’s inequality, we have

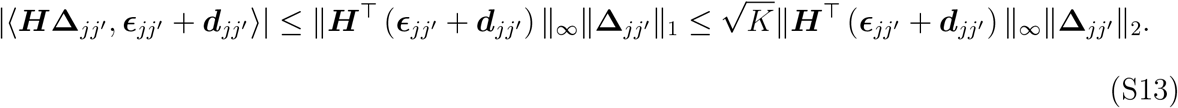

In addition, under Assumption 3, it holds that

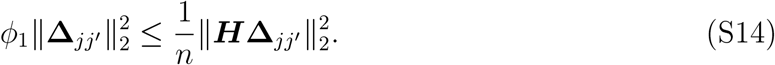

Putting (S12)-(S14) together, we arrive at that

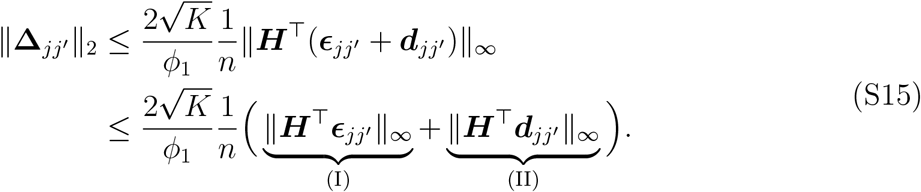

Next, we bound terms (I) and (II) separately. First, we focus on term (I) and bound it with Lemma A3.2. By the union sum inequality, we have

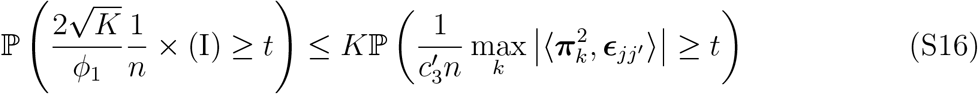

where 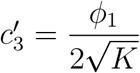.

We note that *ϵ*_*ijj*′_ = *z*_*ij*_*z*_*ij*′_ *−* 𝔼[*z*_*ij*_*z*_*ij*′_]. By Assumption 1, we have 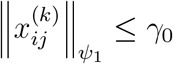 for some constant *γ*_0_ *>* 0, *j* ∈ [*p*], *k* ∈ [*K*]. It follows that 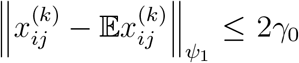. Therefore, there exists a constant *K*_0_ *>* 0 such that 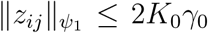 for *j* ∈ [*p*]. Then, let 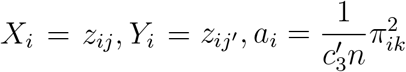 for *i* ∈ [*n*] and γ_*X*_ = γ_*Y*_ = 2*K*_0*γ*0_ in Lemma A3.2, we have

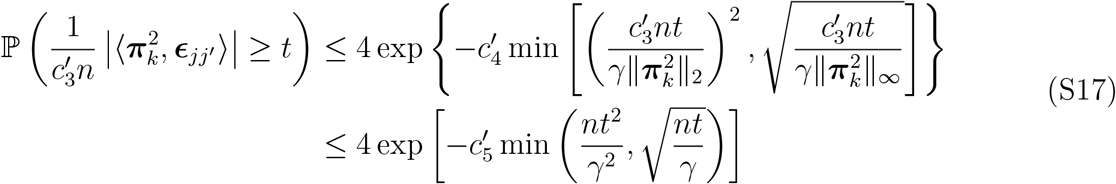

for *k* ∈ [*K*] and some positive constants 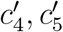, where the last inequality is implied by 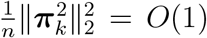 (Assumption 3) and *γ*= (2*K*_0*γ*0_)^2^. Note both cases in Lemma A3.2 are considered to give the result in (S17).

Next, we establish a bound for the second term (II) in (S15). Let 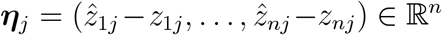 for *j* ∈ [*p*]. Then, we have 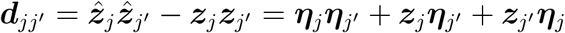, and

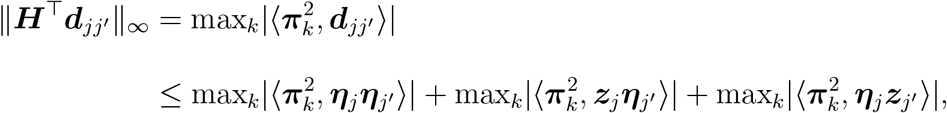

which can be bounded with the following two inequalities:

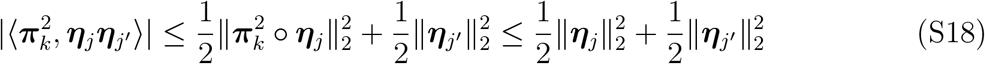

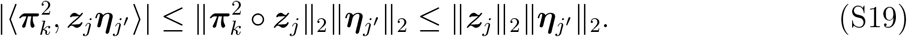

Next, we bound the two terms *‖****η***_*j*_*‖*_2_ and *‖****z***_*j*_*‖*_2_, respectively. For *‖****η***_*j*_*‖*_2_, we have 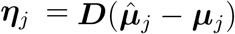, where 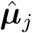 is as defined in (5). By Assumption 3 and using similar arguments as in (S12)-(S14), we have

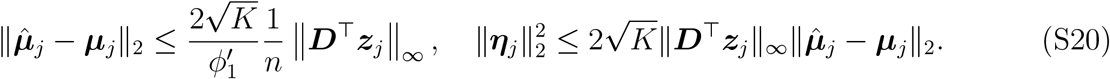

By Hölder’s inequality, we can further combine the equations in (S20) and get

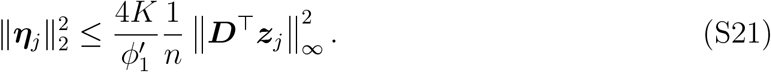

Next, we rewrite term (II) in (S15) as

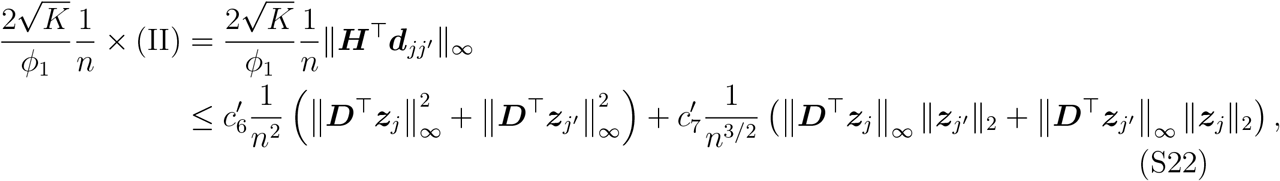

where 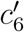 and 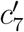 are some positive constants.

To bound 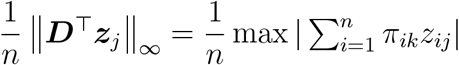, we note that *z*_*ij*_’s are sub-exponential with *‖z*_*ij*_*‖ ≤* 2*K*_0*γ*0_. Therefore, let *a*_*i*_ = *π*_*ik*_, *X*_*i*_ = *z*_*ij*_ for *i* ∈ [*n*] and γ = (2*K*_0*γ*0_)^2^, Lemma A3.1 implies

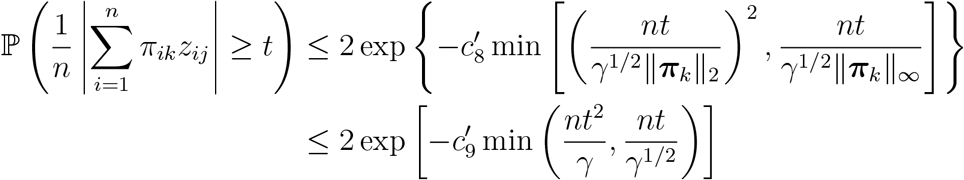

for *k* ∈ [*K*] and some positive constants 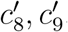, where the last inequality is implied by

Assumption 3 (i.e., 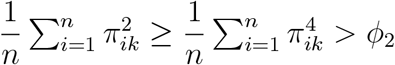). Then, we have

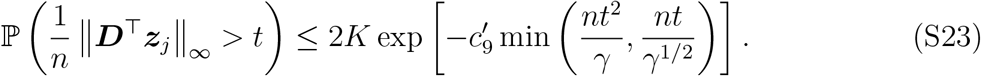

By Lemma A3.5, we have the following result for 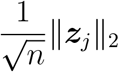:

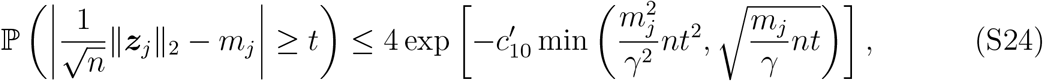

where 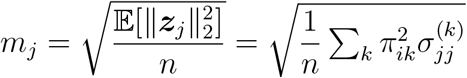 and 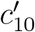 is a positive constant.

Putting (S17) for term (I) and (S23), (S24) for term (II) together, we have

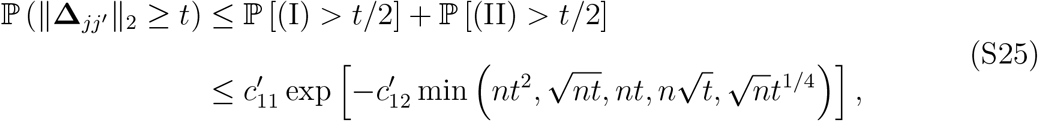

for some positive constants 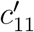 and 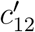. Finally, as 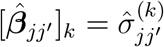 and 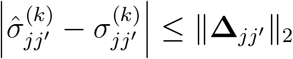, setting 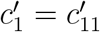 and 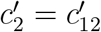, we achieve the desired result in (S10).

**Step 2**. Recall 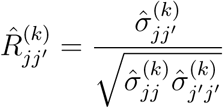. In this step, we show

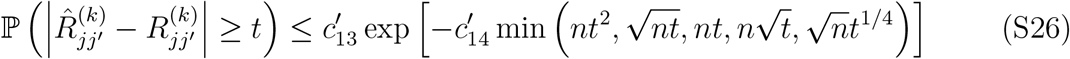

for *k* ∈ [*K*], *j, j*′ ∈ [*p*] and some positive constants 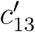 and 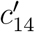. The proof in this step is motivated by the techniques in Jiang (2013).

We first demonstrate that if 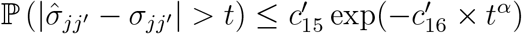 for *j, j*′ ∈ [*p*], some positive constants 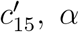, and some positive scalar 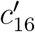 that may depend on *n*. then the corresponding correlation estimate satisfies

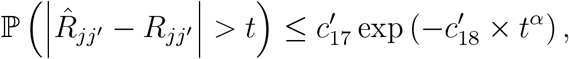

where 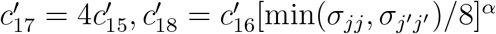. Note that the superscript (*k*) is dropped here and in the ensuing arguments in Step 2, while the derivation holds for all 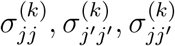 for *k* ∈ [*K*].

Let 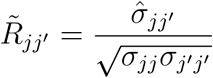 and 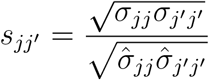. For 0 *<t <* 2, we have

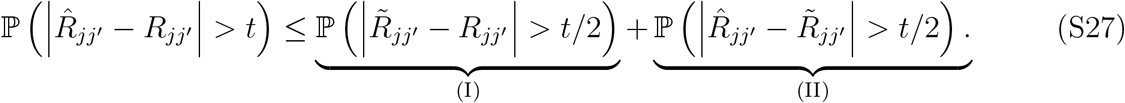

Assuming *σ* _*jj*_, *σ* _*j* ′_ *j* _′_ *>* 0, we have

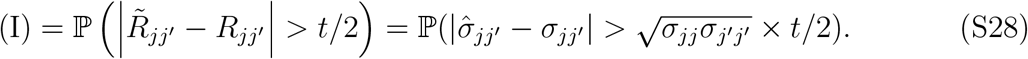

For term (II) in (S27), consider

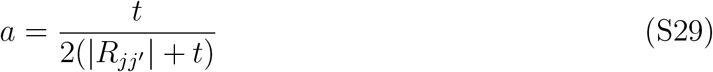

and

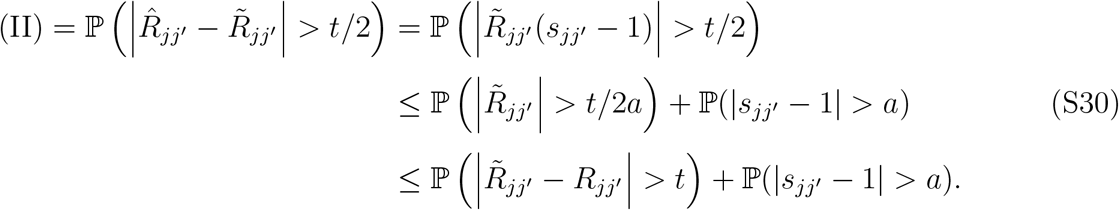

The term 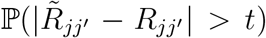 can again be bounded using (S28). Note that *a <* 1, the following holds for the second term

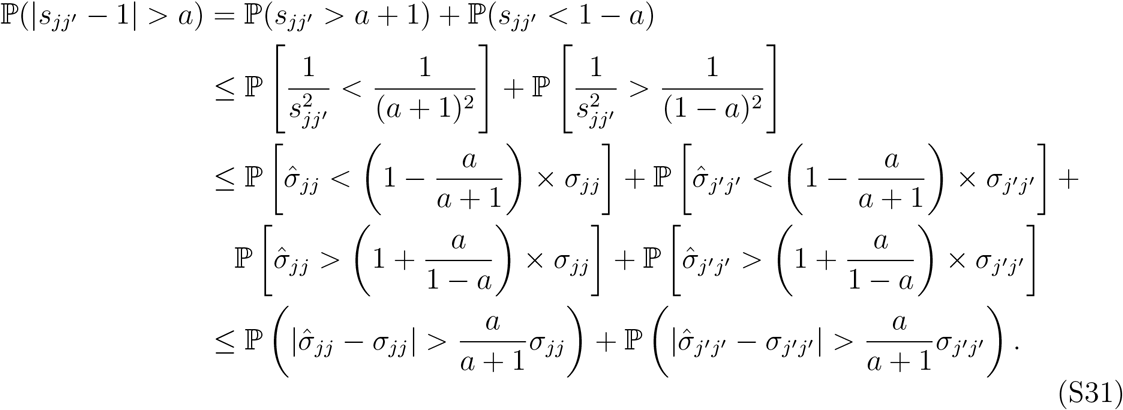

Finally, by (S29) and the fact that *t* ∈ (0, 2), we have 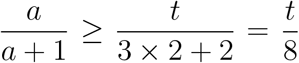. Therefore, summarizing the above terms, we have

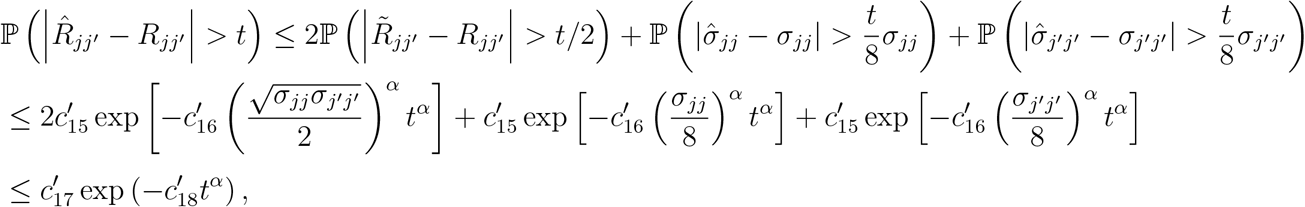

where 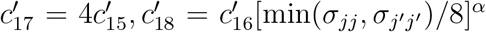. These arguments, combined with the results from Step 1, can be used to establish (S26).

**Step 3**. By Steps 1 and 2, we can then bound the max norm of 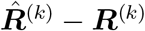 as

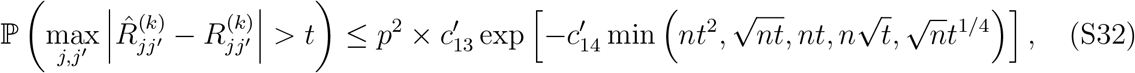

where 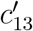 and 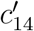 are some positive constants. Suppose log *p* = *o*(*n*^1/3^) and set 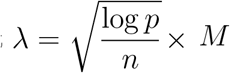 for some large constant *M*. Then, letting *t* = *λ*, we have min 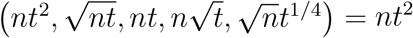 and (S32) can be written as

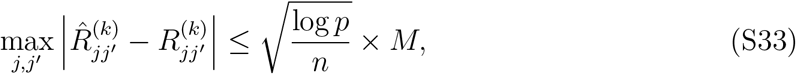

holds with probability at least 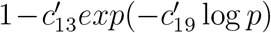, where 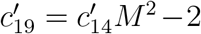 is a large positive constant. Combined with Lemma A3.4, we have

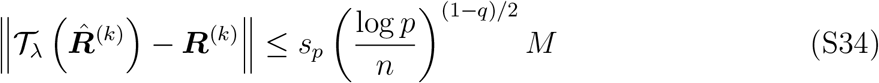

with probability at least 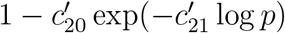 for some positive constants 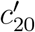 and 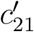.

#### A3.3 Proof of Theorem 5.2

As it holds that

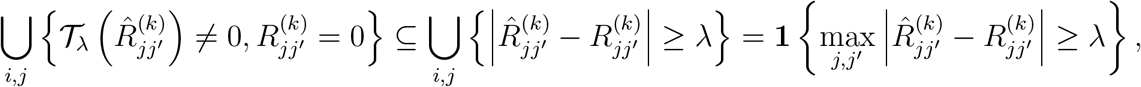

we have by (S32), log *p* = *o*(*n*^1*/*3^) and 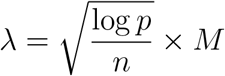 for some large constant *M* that

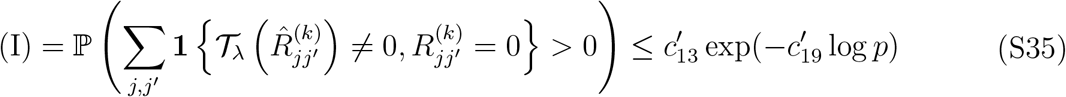

for 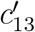 and 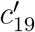 as defined in Section A3.2.

Given 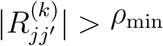 and 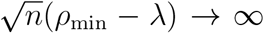, we have

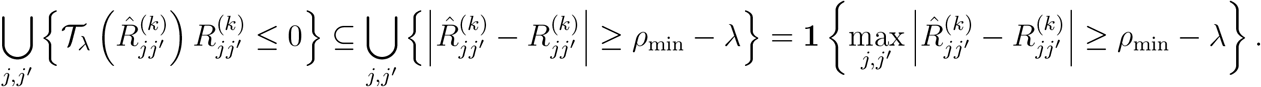

Combined with (S32), we obtain

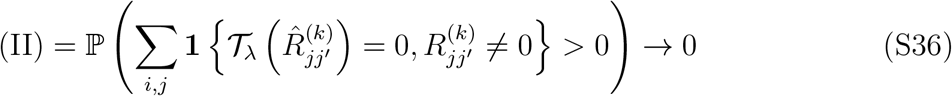

As 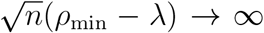. Finally, as 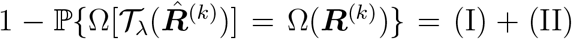, we obtain 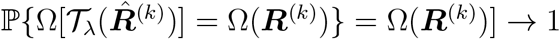.

#### A3.4 Proof of Lemma A3.2

The proof is divided into two steps. In step 1, we will establish the tail bound for the weighted sum of fourth degree polynomials of sub-Gaussian random variables. Next, in step 2, we will apply the established tail bound to product of sub-exponential random variables by representing centered products of sub-exponential random variables as a fourth degree polynomial of sub-Gaussian random variables.

**Step 1:** Following Adamczak and Wolff (2015), we define a norm for tensors that will be used in our concentration result. Let *d* ∈ ℕ _+_ be a positive integer. We denote by *𝒫*_*d*_ the set of its partitions of [*d*] into non-empty and non-intersecting disjoint sets. Moreover, let 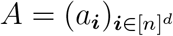 be a tensor of order-*d*, whose entries are of the form

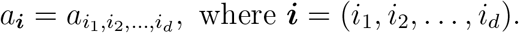

Let *𝒥* = {*J*_1_, …, *J*_*k*_} ∈ *𝒫* _*d*_ be a fixed partition of [*d*], where *J*_*j*_ ⊆ [*d*] for each *j* ∈ [*k*]. Let | *𝒥* | denote the cardinality of the *𝒥*, which is equal to *k*. We define a norm ‖·‖_*𝒥*_ by

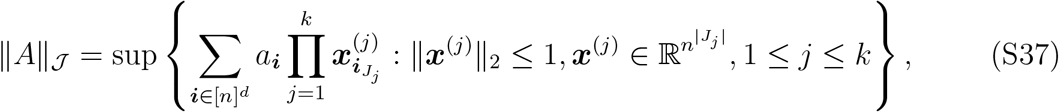

where we write ***i***_*I*_ = (*i*_*k*_)_*k*∈*I*_ for any *I* ⊆ [*d*] and the supremum is taken over all possible *k* vectors {***x***^(1)^, …, ***x***^(*k*)^}.

Let *f* (*u*) = *u*^4^ and consider a polynomial of degree 4:

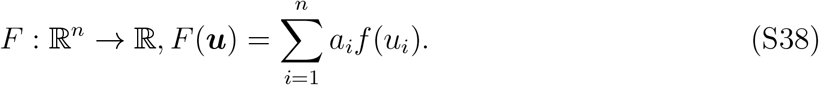

Let ***Z*** = [*Z*_1_, …, *Z*_*n*_]^T^ be a vector of independent components with 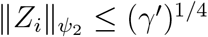, where *γ*′ is some positive constant. Plugging in *F* (*Z*), letting *D* = 4, Lemma A3.3 gives

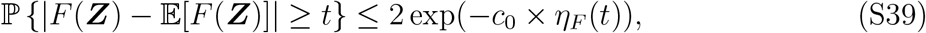

and

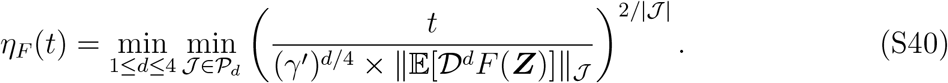

Following Lemma 3 of Balasubramanian et al. (2018), we have

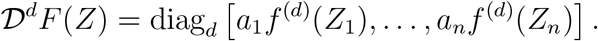

Using (31) in Balasubramanian et al. (2018) and the fact that |*f* ^(*d*)^(*u*)| *≤* 4!|*u*|^4− *d*^ for *d* = 1, …, 4, we obtain

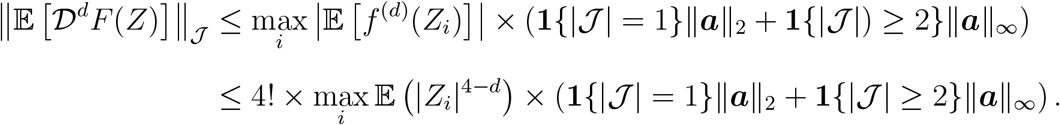

By the definition of *ψ*_2_-norm, 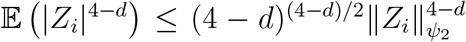. In addition, recall that 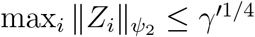, then

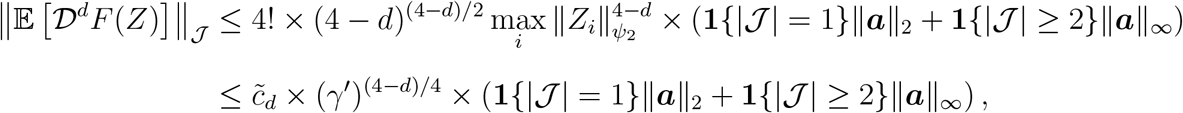

where 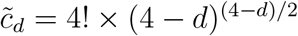 for *d* = 1, …, 4. Then,

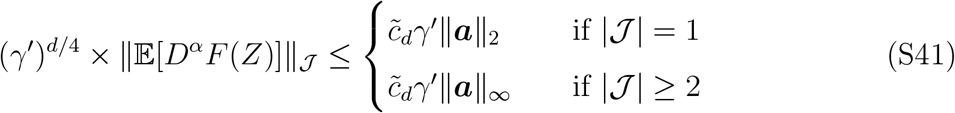

Combining (S40) and (S41), we have

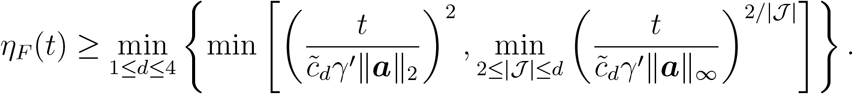

If *t< c*_min_ *× γ*′ *× ‖****a****‖* _*∞*_ for 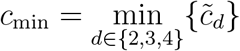, then

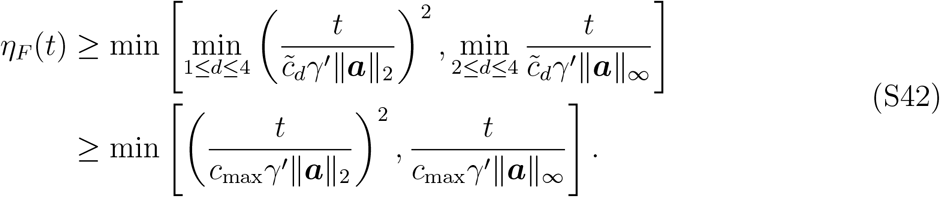

where 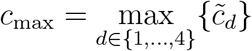. Plugging it in Lemma A3.3, we obtain

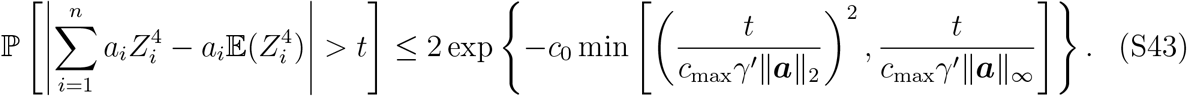

If *t ≥ c*_max_ *× γ*′ *× ‖****a****‖* _*∞*_, then

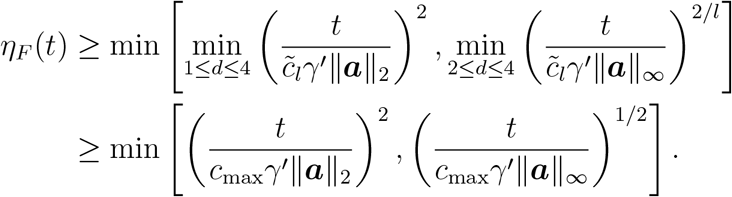

Similarly, we obtain

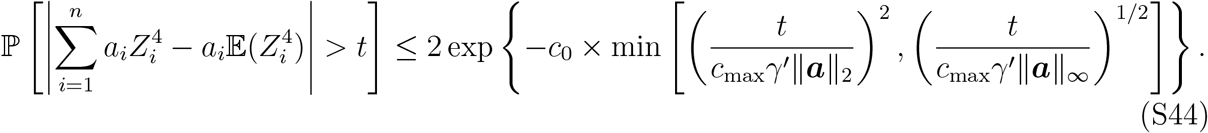

**Step 2**: Next, we show how a product of sub-exponential random variables can be represented as a fourth degree polynomial of Gaussian random variables, and then leverage the bound established in step 1 here.

Let 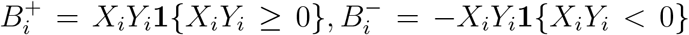. It follows that 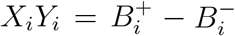. Furthermore, define 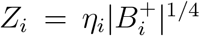 for *i* = 1, …, *n*, where 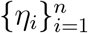 are independent Rademacher random variables. Then, we have

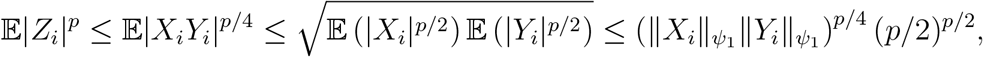

where the second inequality is given by Cauchy-Schwarz inequality and the third is given by the definition of _1_-norm in Assumption 1. Then, by the definition of *ψ*_2_-norm,

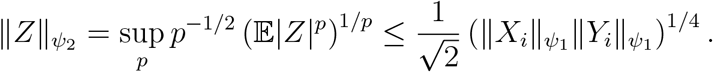

Therefore, for *i* = 1, …, *n, Z*_*i*_ are sub-Gaussian random variables with

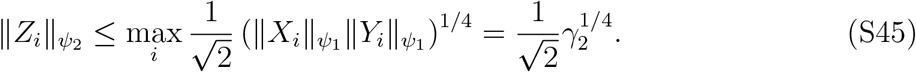

Next, we start to apply the results obtained in step 1. First consider the case where 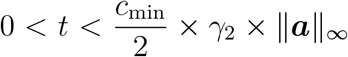. Then, plugging *γ*′ = *γ*_2_*/*4 and the tail cutoff *t/*2 into (S43), we obtain

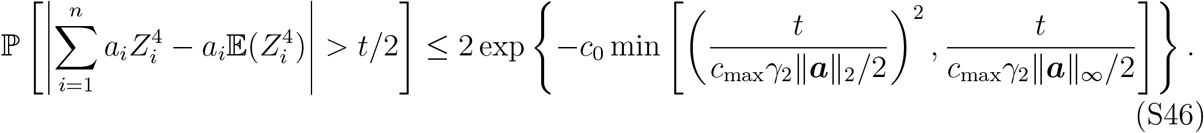

As 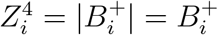, it follows that

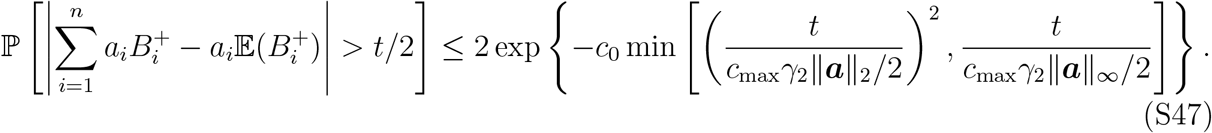

By symmetry, similar derivations give the same bound for 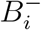as (S47). Then

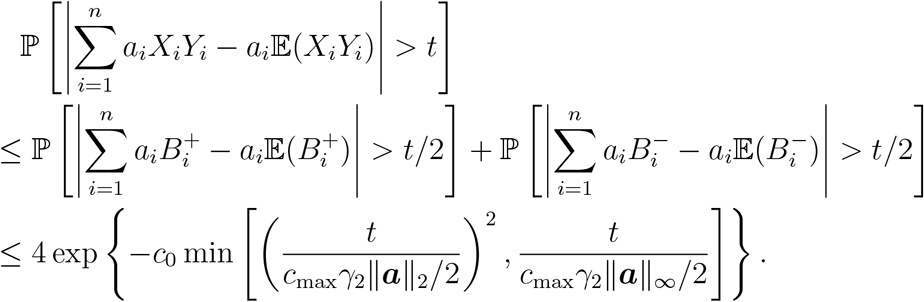

Finally, consider the case where 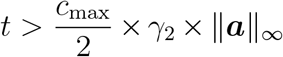. Following a similar argument, we get

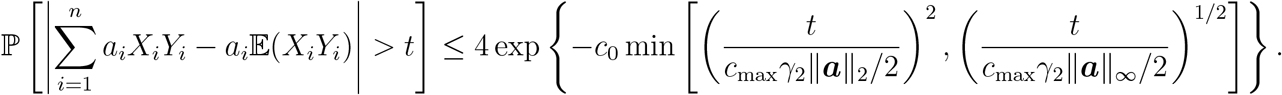

Then, setting 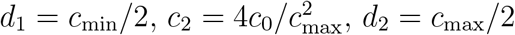 gives the desired inequalities.

#### A3.5 Proof of Lemma A3.5

Given 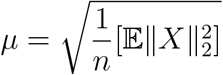, by Lemma A3.2, we have

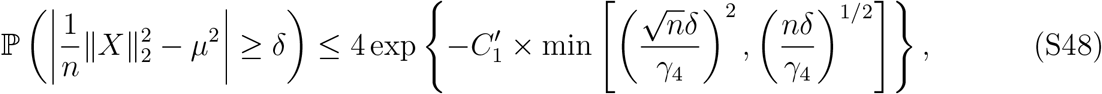

for any positive constant *δ* and some positive constant 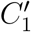. Equivalently, let 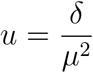, then

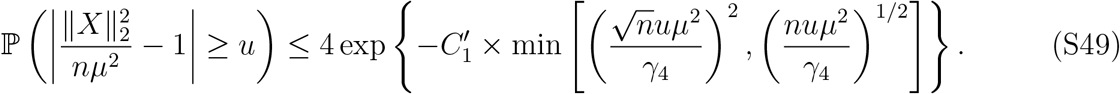

As |*z −* 1| *>u* implies |*z*^2^ *−* 1| *>* max(*u, u*^2^) for any *z ≥* 0. Then, if *u ≤* 1,

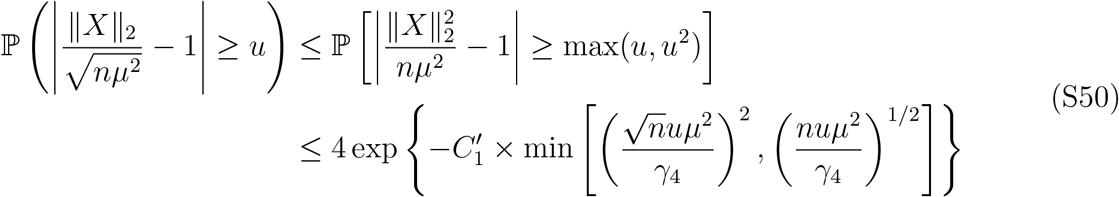

Setting *t* = *uμ* and 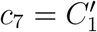 gives (S9).

#### A3.6 Extended discussions on Assumptions 3

Recall

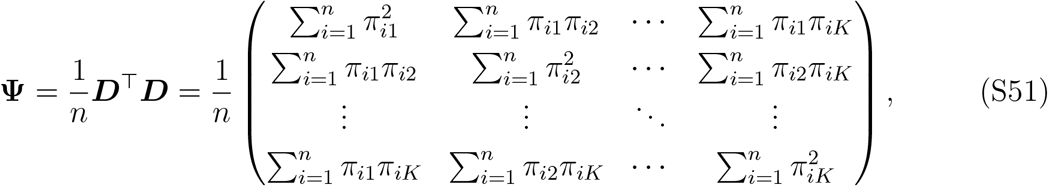

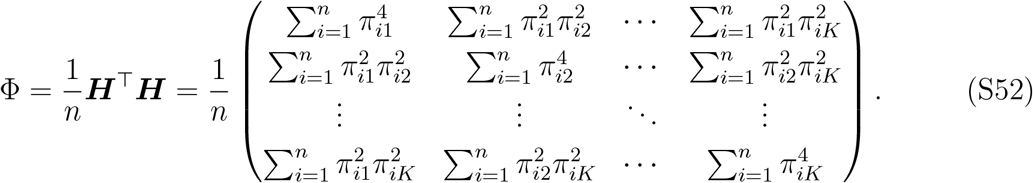

First, suppose (*π*_*i*1_, …, *π*_*iK*_) ∼ Multinomial(1, ***θ***), where ***θ*** = (*θ*_1_, …, *θ*_*K*_) ∈ ℝ^*K*^ is a vector of positive parameters summed to 1. Though this assumption on cell type proportions implies that each tissue sample can have cells from only one of the *K* cell types, it nevertheless provides some insight into (S51) and (S52). Specifically, the smallest eigenvalue of

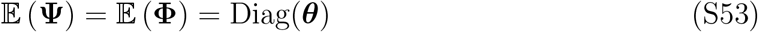

is min{*θ*_*k*_}, a positive constant that does not depend on *n*.

Next, consider a less simplified model with (*π*_*i*1_, …, *π*_*iK*_) ∼ Dirichlet(***α***, where ***α*** = (*α*_1_, …, *α*_*K*_) ∈ ℝ^*K*^ is a vector of postive parameters. Let *α*_0_ = Σ _*k*_ *α*_*k*_. Then,

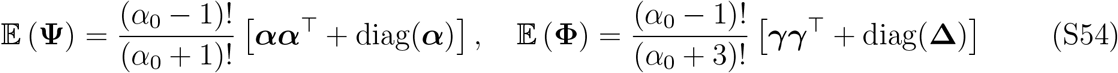

where γ = (***α*** + 1) ° ***α*** ∈ ℝ^*K*^, Δ= ***α*** ° (***α*** + 1) ° (4***α***+ 6) ∈ ℝ^*K*^. The smallest eigenvalue of 𝔼(Ψ) and 𝔼(Φ) are lower bounded by positive constants, 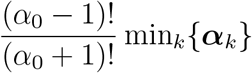 and 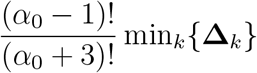, respectively.

The above derivation demonstrates that under reasonable probabilistic models, the eigen-value conditions in Assumption 3 are satisfied at the population level. This bounded norm condition in Assumption 3 requires the sum of the squared proportions to grow linearly with the size of the population. This is plausible when considering any common cell types: a cell type is defined as “common” if 𝔼[*π*_*ik*_] *> C*_*k*_ for some constant *C*_*k*_ *>* 0, which also implies 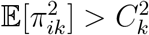.

**Figure S9:**
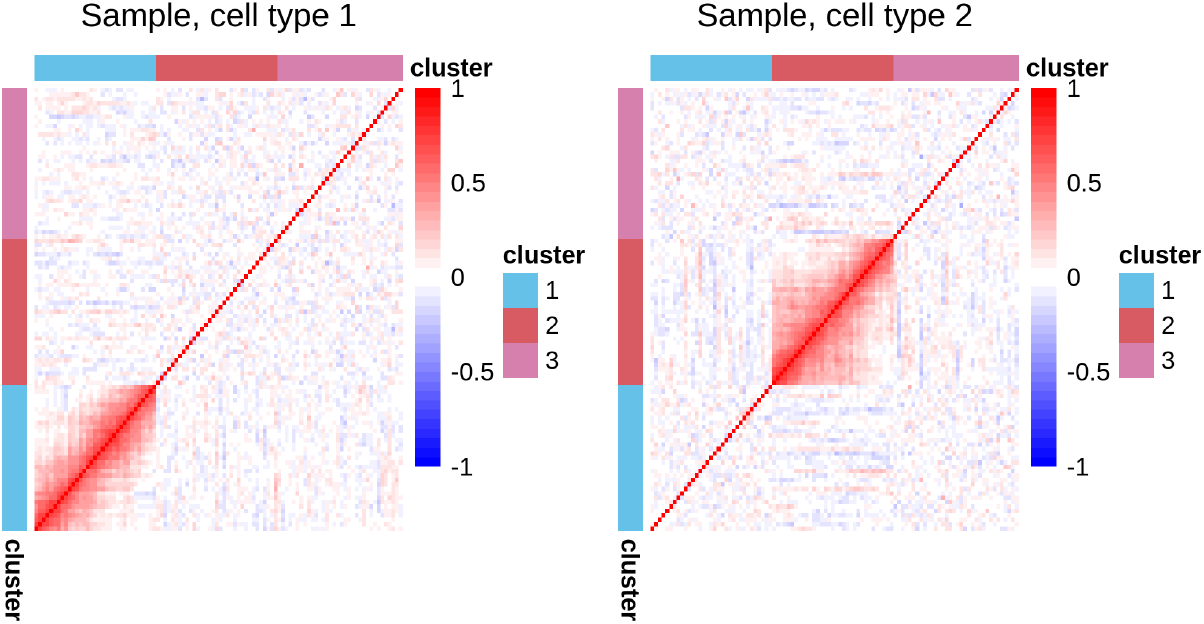
Sample correlations from *n* = 150 samples of *p* = 100 genes, with data simulated from Algorithm 1 under a multivariate negative binomial distribution with an AR(1)-type correlation structure in *V*_1_ for cell type 1 and *V*_2_ for cell type 2 (See Section 3).

### A4 More Results from Simulation studies in Section 3

In Section 4, we followed the `bMIND` tutorial at https://htmlpreview.github.io/?https://github.com/randel/MIND/blob/master/bMIND_tutorial.html to estimate cell-type-specific expression levels in each sample using `bMIND`. There are two steps provided by the `bMIND` implementation: first, use single cell data to infer priors for each gene; second, fit the Bayesian mixed effects model to infer posterior means in each sample. However, due to the high dropout rates in single cell data and/or the low expression levels, some genes may have zero UMI counts detected, which is infeasible to infer priors. Therefore, only a subset of genes would have valid priors inferred by `bMIND`. For those genes, we supplied the valid priors as informative priors to `bMIND` for posterior inference, as recommended by the tutorial. For the other genes, we used the default non-informative priors in `bMIND`.

**Table S1:** Evaluation criteria under the same setting as Table 1. `bMIND (inf)` denotes the co-expression estimates computed with `bMIND` given informative priors. We use F-norm to denote the Frobenius norm and Op.-norm to denote the operator norm.

| | | | Cell type 1 ($m = 2/3$) | | | | Cell type 2 ($m = 1/3$) | | | |
| --- | --- | --- | --- | --- | --- | --- | --- | --- | --- | --- |
| $n$ | $p$ | Method | F-norm | Op.-norm | FPR | TPR | F-norm | Op.-norm | FPR | TPR |
| 150 | 100 | **bMIND (inf)** | 9.05<br>(0.10) | 2.54<br>(0.12) | - | - | 11.76<br>(0.16) | 3.82<br>(0.23) | - | - |
|  | 200 | **bMIND (inf)** | 18.02<br>(0.10) | 4.20<br>(0.15) | - | - | 23.41<br>(0.19) | 6.70<br>(0.35) | - | - |
| 600 | 100 | **bMIND (inf)** | 4.72<br>(0.05) | 1.29<br>(0.01) | - | - | 6.41<br>(0.08) | 1.77<br>(0.12) | - | - |
|  | 200 | **bMIND (inf)** | 9.17<br>(0.05) | 1.72<br>(0.05) | - | - | 12.53<br>(0.09) | 3.15<br>(0.20) | - | - |

**Table S2:** Evaluation criteria with varying sample size *n* and network size *p* under a negative binomial distribution with an AR(1)-type correlation structure in *V*_1_ or *V*_2_ (See Section 3). The four methods under comparison are `Bulk`, `d-CSNet`, `CSNet`, and `bMIND`. We use F-norm to denote the Frobenius norm and Op.-norm to denote the operator norm. Marked in boldface are those achieving the best evaluation criteria in each setting.

| | | | Cell type 1 ($m = 2/3$) | | | | Cell type 2 ($m = 1/3$) | | | |
| --- | --- | --- | --- | --- | --- | --- | --- | --- | --- | --- |
| $n$ | $p$ | Method | F-norm | Op.-norm | FPR | TPR | F-norm | Op.-norm | FPR | TPR |
| 150 | 100 | Bulk | 25.72<br>(0.21) | 24.67<br>(0.17) | 0.32<br>(0.05) | **0.75**<br>(**0.06**) | 28.55<br>(0.26) | 24.67<br>(0.17) | 0.36<br>(0.04) | **0.43**<br>(**0.07**) |
|  |  | d-CSNet | 11.82<br>(0.28) | 4.78<br>(0.63) | -<br>- | -<br>- | 25.87<br>(0.98) | 10.13<br>(1.08) | -<br>- | -<br>- |
|  |  | CSNet | **5.39**<br>(**0.59**) | **3.58**<br>(**0.73**) | **0.16**<br>(**0.04**) | 0.72<br>(0.07) | **10.85**<br>(**0.54**) | **8.43**<br>(**0.69**) | **0.08**<br>(**0.02**) | 0.37<br>(0.09) |
|  |  | bMIND | 12.07<br>(0.26) | 7.22<br>(0.43) | -<br>- | -<br>- | 16.48<br>(0.42) | 9.48<br>(0.19) | -<br>- | -<br>- |
|  | 200 | Bulk | 49.91<br>(0.24) | 48.58<br>(0.21) | 0.26<br>(0.03) | **0.48**<br>(**0.04**) | 52.98<br>(0.23) | 48.58<br>(0.21) | 0.28<br>(0.03) | **0.27**<br>(**0.04**) |
|  |  | d-CSNet | 23.76<br>(0.30) | 8.15<br>(0.74) | -<br>- | -<br>- | 51.50<br>(1.39) | 18.56<br>(1.76) | -<br>- | -<br>- |
|  |  | CSNet | **9.19**<br>(**0.57**) | **5.82**<br>(**0.65**) | **0.11**<br>(**0.02**) | 0.46<br>(0.04) | **17.14**<br>(**0.40**) | **11.41**<br>(**0.49**) | **0.04**<br>(**0.02**) | 0.15<br>(0.05) |
|  |  | bMIND | 20.99<br>(0.27) | 8.97<br>(0.39) | -<br>- | -<br>- | 27.84<br>(0.48) | 11.61<br>(0.37) | -<br>- | -<br>- |
| 600 | 100 | Bulk | 25.54<br>(0.10) | 24.78<br>(0.08) | 0.37<br>(0.05) | **0.90**<br>(**0.04**) | 28.42<br>(0.17) | 24.78<br>(0.08) | 0.40<br>(0.05) | 0.67<br>(0.05) |
|  |  | d-CSNet | 5.82<br>(0.12) | 2.17<br>(0.29) | -<br>- | -<br>- | 12.08<br>(0.28) | 4.20<br>(0.53) | -<br>- | -<br>- |
|  |  | CSNet | **2.73**<br>(**0.34**) | **1.63**<br>(**0.42**) | **0.13**<br>(**0.07**) | 0.86<br>(0.05) | **5.72**<br>(**0.53**) | **3.68**<br>(**0.77**) | **0.16**<br>(**0.05**) | **0.71**<br>(**0.07**) |
|  |  | bMIND | 8.89<br>(0.10) | 6.77<br>(0.18) | -<br>- | -<br>- | 12.53<br>(0.31) | 8.82<br>(0.21) | -<br>- | -<br>- |
|  | 200 | Bulk | 49.66<br>(0.13) | 48.79<br>(0.12) | 0.30<br>(0.03) | **0.63**<br>(**0.04**) | 53.03<br>(0.17) | 48.79<br>(0.12) | 0.32<br>(0.03) | **0.43**<br>(**0.05**) |
|  |  | d-CSNet | 11.73<br>(0.14) | 3.54<br>(0.37) | -<br>- | -<br>- | 24.18<br>(0.37) | 7.19<br>(0.71) | -<br>- | -<br>- |
|  |  | CSNet | **4.79**<br>(**0.32**) | **2.82**<br>(**0.49**) | **0.10**<br>(**0.04**) | 0.57<br>(0.05) | **9.71**<br>(**0.54**) | **6.10**<br>(**0.71**) | **0.11**<br>(**0.03**) | **0.43**<br>(**0.04**) |
|  |  | bMIND | 13.42<br>(0.11) | 8.42<br>(0.21) | -<br>- | -<br>- | 19.58<br>(0.40) | 10.46<br>(0.24) | -<br>- | -<br>- |

**Table S3:**
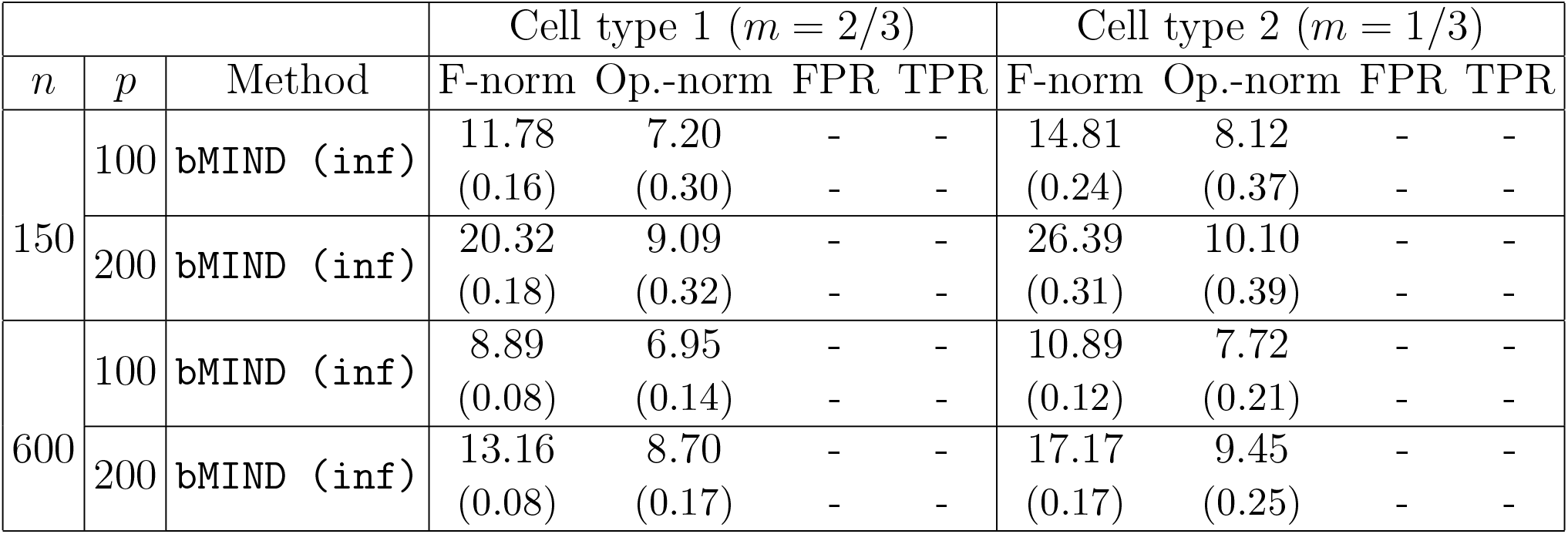
Evaluation criteria under the same setting as Table S2. `bMIND (inf)` denotes the co-expression estimates computed with `bMIND` given informative priors. We use F-norm to denote the Frobenius norm and Op.-norm to denote the operator norm.

## References

Abbas, A. R., Wolslegel, K., Seshasayee, D., Modrusan, Z., and Clark, H. F. (2009), “Deconvolution of blood microarray data identifies cellular activation patterns in systemic lupus erythematosus,” PloS one, 4, e6098.

Adamczak, R. and Wol?, P. (2015), “Concentration inequalities for non-Lipschitz functions with bounded derivatives of higher order,” Probability Theory and Related Fields, 162, 531–586.

Alzheimer’s Association (2019), “2019 Alzheimer’s disease facts and figures,” Alzheimer’s & dementia, 15, 321–387.

Ashburner, M., Ball, C. A., Blake, J. A., Botstein, D., Butler, H., Cherry, J. M., Davis, A. P., Dolinski, K., Dwight, S. S., Eppig, J. T., et al. (2000), “Gene ontology: tool for the unification of biology,” Nature genetics, 25, 25–29.

Balasubramanian, K., Fan, J., and Yang, Z. (2018), “Tensor methods for additive index models under discordance and heterogeneity,” arXiv preprint 1807.06693.

Bennett, D. A., Buchman, A. S., Boyle, P. A., Barnes, L. L., Wilson, R. S., and Schneider, J. A. (2018), “Religious orders study and rush memory and aging project,” Journal of Alzheimer’s disease, 64, S161–S189.

Bickel, P. J. and Levina, E. (2008a), “Covariance regularization by thresholding,” The Annals of Statistics, 36, 2577–2604.

Bickel, P. J. and Levina, E. (2008b), “Regularized estimation of large covariance matrices,” The Annals of Statistics, 36, 199–227.

Cai, Z. and Xiao, M. (2016), “Oligodendrocytes and Alzheimer’s disease,” International Journal of Neuroscience, 126, 97–104.

Chen, Y., Lun, A. T., and Smyth, G. K. (2014), “Differential expression analysis of complex RNA-seq experiments using edgeR,” Statistical analysis of next generation sequencing data, 51–74.

Cobos, F. A., Alquicira-Hernandez, J., Powell, J. E., Mestdagh, P., and De Preter, K. (2020), “Benchmarking of cell type deconvolution pipelines for transcriptomics data,” Nature communications, 11, 1–14.

Consortium, T. G. O. (2021), “The Gene Ontology resource: enriching a GOld mine,” Nucleic Acids Research, 49, D325–D334.

Deming, Y., Filipello, F., Cignarella, F., Cantoni, C., Hsu, S., Mikesell, R., Li, Z., Del-Aguila, J. L., Dube, U., Farias, F. G., et al. (2019), “The MS4A gene cluster is a key modulator of soluble TREM2 and Alzheimer’s disease risk,” Science translational medicine, 11.

Denisenko, E., Guo, B. B., Jones, M., Hou, R., De Kock, L., Lassmann, T., Poppe, D., Clément, O., Simmons, R. K., Lister, R., et al. (2020), “Systematic assessment of tissue dissociation and storage biases in single-cell and single-nucleus RNA-seq workflows,” Genome biology, 21, 1–25.

Dong, M., Thennavan, A., Urrutia, E., Li, Y., Perou, C. M., Zou, F., and Jiang, Y. (2021), “SCDC: bulk gene expression deconvolution by multiple single-cell RNA sequencing references,” Briefings in bioinformatics, 22, 416–427.

El Karoui, N. (2008), “Operator norm consistent estimation of large-dimensional sparse covariance matrices,” The Annals of Statistics, 36, 2717–2756.

Fan, J. and Li, R. (2001), “Variable selection via nonconcave penalized likelihood and its oracle properties,” Journal of the American statistical Association, 96, 1348–1360.

Gaiteri, C., Ding, Y., French, B., Tseng, G. C., and Sibille, E. (2014), “Beyond modules and hubs: the potential of gene coexpression networks for investigating molecular mechanisms of complex brain disorders,” Genes, brain and behavior, 13, 13–24.

Griciuc, A., Serrano-Pozo, A., Parrado, A. R., Lesinski, A. N., Asselin, C. N., Mullin, K., Hooli, B., Choi, S. H., Hyman, B. T., and Tanzi, R. E. (2013), “Alzheimer’s disease risk gene CD33 inhibits microglial uptake of amyloid beta,” Neuron, 78, 631–643.

Heintzman, N. D., Hon, G. C., Hawkins, R. D., Kheradpour, P., Stark, A., Harp, L. F., Ye, Z., Lee, L. K., Stuart, R. K., Ching, C. W., et al. (2009), “Histone modifications at human enhancers reflect global cell-type-specific gene expression,” Nature, 459, 108–112.

Heneka, M. T., Carson, M. J., El Khoury, J., Landreth, G. E., Brosseron, F., Feinstein, D. L., Jacobs, A. H., Wyss-Coray, T., Vitorica, J., Ransoho?, R. M., et al. (2015), “Neuroinflammation in Alzheimer’s disease,” The Lancet Neurology, 14, 388–405.

Higham, N. J. (2002), “Computing the nearest correlation matrix—a problem from finance,” IMA journal of Numerical Analysis, 22, 329–343.

Hwang, B., Lee, J. H., and Bang, D. (2018), “Single-cell RNA sequencing technologies and bioinformatics pipelines,” Experimental & molecular medicine, 50, 1–14.

Jew, B., Alvarez, M., Rahmani, E., Miao, Z., Ko, A., Garske, K. M., Sul, J. H., Pietiläinen, K. H., Pajukanta, P., and Halperin, E. (2020), “Accurate estimation of cell composition in bulk expression through robust integration of single-cell information,” Nature communications, 11, 1–11.

Jiang, B. (2013), “Covariance selection by thresholding the sample correlation matrix,” Statistics & Probability Letters, 83, 2492–2498.

Kiselev, V. Y., Andrews, T. S., and Hemberg, M. (2019), “Challenges in unsupervised clustering of single-cell RNA-seq data,” Nature Reviews Genetics, 20, 273–282.

Kosoy, R., Fullard, J., Zeng, B., Bendl, J., Dong, P., Rahman, S., Kleopoulos, S., Shao, Z., Humphrey, J., de Paiva Lopes, K., et al. (2021), “Genetics of the human microglia regulome refines Alzheimer’s disease risk loci,” medRxiv.

Langfelder, P. and Horvath, S. (2008), “WGCNA: an R package for weighted correlation network analysis,” BMC bioinformatics, 9, 1–13.

Li, B., Severson, E., Pignon, J.-C., Zhao, H., Li, T., Novak, J., Jiang, P., Shen, H., Aster, J. C., Rodig, S., et al. (2016), “Comprehensive analyses of tumor immunity: implications for cancer immunotherapy,” Genome biology, 17, 1–16.

Love, M. I., Huber, W., and Anders, S. (2014), “Moderated estimation of fold change and dispersion for RNA-seq data with DESeq2,” Genome biology, 15, 1–21.

Mathys, H., Davila-Velderrain, J., Peng, Z., Gao, F., Mohammadi, S., Young, J. Z., Menon, M., He, L., Abdurrob, F., Jiang, X., et al. (2019), “Single-cell transcriptomic analysis of Alzheimer’s disease,” Nature, 570, 332–337.

Meng, G. and Mei, H. (2019), “Transcriptional dysregulation study reveals a core network involving the progression of Alzheimer’s disease,” Frontiers in aging neuroscience, 11, 101.

Morabito, S., Miyoshi, E., Michael, N., Shahin, S., Martini, A. C., Head, E., Silva, J., Leavy, K., Perez-Rosendahl, M., and Swarup, V. (2021), “Single-nucleus chromatin accessibility and transcriptomic characterization of Alzheimer’s disease,” Nature Genetics, 53, 1143– 1155.

Mostafavi, S., Gaiteri, C., Sullivan, S. E., White, C. C., Tasaki, S., Xu, J., Taga, M., Klein, H.-U., Patrick, E., Komashko, V., et al. (2018), “A molecular network of the aging human brain provides insights into the pathology and cognitive decline of Alzheimer’s disease,” Nature neuroscience, 21, 811–819.

Newman, A. M., Liu, C. L., Green, M. R., Gentles, A. J., Feng, W., Xu, Y., Hoang, C. D., Diehn, M., and Alizadeh, A. A. (2015), “Robust enumeration of cell subsets from tissue expression profiles,” Nature methods, 12, 453–457.

Newman, A. M., Steen, C. B., Liu, C. L., Gentles, A. J., Chaudhuri, A. A., Scherer, F., Khodadoust, M. S., Esfahani, M. S., Luca, B. A., Steiner, D., et al. (2019), “Determining cell type abundance and expression from bulk tissues with digital cytometry,” Nature biotechnology, 37, 773–782.

Pimenova, A. A., Raj, T., and Goate, A. M. (2018), “Untangling genetic risk for Alzheimer’s disease,” Biological psychiatry, 83, 300–310.

Robinson, M. D., McCarthy, D. J., and Smyth, G. K. (2010), “edgeR: a Bioconductor package for differential expression analysis of digital gene expression data,” Bioinformatics, 26, 139–140.

Ronning, G. (1977), “A simple scheme for generating multivariate gamma distributions with non-negative covariance matrix,” Technometrics, 19, 179–183.

Rothman, A. J., Bickel, P. J., Levina, E., and Zhu, J. (2008), “Sparse permutation invariant covariance estimation,” Electronic Journal of Statistics, 2, 494–515.

Rothman, A. J., Levina, E., and Zhu, J. (2009), “Generalized thresholding of large covariance matrices,” Journal of the American Statistical Association, 104, 177–186.

Sims, R., Hill, M., and Williams, J. (2020), “The multiplex model of the genetics of Alzheimer’s disease,” Nature neuroscience, 23, 311–322.

Sims, R., Van Der Lee, S. J., Naj, A. C., Bellenguez, C., Badarinarayan, N., Jakobsdottir, J., Kunkle, B. W., Boland, A., Raybould, R., Bis, J. C., et al. (2017), “Rare coding variants in PLCG2, ABI3, and TREM2 implicate microglial-mediated innate immunity in Alzheimer’s disease,” Nature genetics, 49, 1373–1384.

Skene, N. G. and Grant, S. G. (2016), “Identification of vulnerable cell types in major brain disorders using single cell transcriptomes and expression weighted cell type enrichment,” Frontiers in neuroscience, 10, 16.

Stower, H. (2019), “Single-cell insights into neurology,” Nature medicine, 25, 1799–1799.

Tang, D., Park, S., and Zhao, H. (2020), “NITUMID: nonnegative matrix factorization-based immune-TUmor MIcroenvironment Deconvolution,” Bioinformatics, 36, 1344–1350.

Tansey, K. E., Cameron, D., and Hill, M. J. (2018), “Genetic risk for Alzheimer’s disease is concentrated in specific macrophage and microglial transcriptional networks,” Genome medicine, 10, 1–10.

Trapnell, C., Williams, B. A., Pertea, G., Mortazavi, A., Kwan, G., Van Baren, M. J., Salzberg, S. L., Wold, B. J., and Pachter, L. (2010), “Transcript assembly and quantification by RNA-Seq reveals unannotated transcripts and isoform switching during cell differentiation,” Nature biotechnology, 28, 511–515.

Vershynin, R. (2018), High-dimensional probability: An introduction with applications in data science, vol. 47, Cambridge university press.

Wan, Y.-W., Al-Ouran, R., Mangleburg, C. G., Perumal, T. M., Lee, T. V., Allison, K., Swarup, V., Funk, C. C., Gaiteri, C., Allen, M., et al. (2020), “Meta-analysis of the Alzheimer’s disease human brain transcriptome and functional dissection in mouse models,” Cell reports, 32, 107908.

Wang, J., Roeder, K., and Devlin, B. (2021a), “Bayesian estimation of cell type-specific gene expression with prior derived from single-cell data,” Genome Research, gr–268722.

Wang, M., Li, A., Sekiya, M., Beckmann, N. D., Quan, X., Schrode, N., Fernando, M. B., Yu, A., Zhu, L., Cao, J., et al. (2021b), “Transformative network modeling of multi-omics data reveals detailed circuits, key regulators, and potential therapeutics for Alzheimer’s disease,” Neuron, 109, 257–272.

Wang, M., Roussos, P., McKenzie, A., Zhou, X., Kajiwara, Y., Brennand, K. J., De Luca, G. C., Crary, J. F., Casaccia, P., Buxbaum, J. D., et al. (2016), “Integrative network analysis of nineteen brain regions identifies molecular signatures and networks underlying selective regional vulnerability to Alzheimer’s disease,” Genome medicine, 8, 1–21.

Wang, X., Park, J., Susztak, K., Zhang, N. R., and Li, M. (2019), “Bulk tissue cell type deconvolution with multi-subject single-cell expression reference,” Nature communications, 10, 1–9.

Winblad, B., Amouyel, P., Andrieu, S., Ballard, C., Brayne, C., Brodaty, H., CedazoMinguez, A., Dubois, B., Edvardsson, D., Feldman, H., et al. (2016), “Defeating Alzheimer’s disease and other dementias: a priority for European science and society,” The Lancet Neurology, 15, 455–532.

Yamazaki, Y., Zhao, N., Caulfield, T. R., Liu, C.-C., and Bu, G. (2019), “Apolipoprotein E and Alzheimer disease: pathobiology and targeting strategies,” Nature Reviews Neurology, 15, 501–518.

Yang, T., Alessandri-Haber, N., Fury, W., Schaner, M., Breese, R., LaCroix-Fralish, M., Kim, J., Adler, C., Macdonald, L. E., Atwal, G. S., et al. (2021), “AdRoit is an accurate and robust method to infer complex transcriptome composition,” Communications Biology, 4, 1–14.

Zhang, B., Gaiteri, C., Bodea, L.-G., Wang, Z., McElwee, J., Podtelezhnikov, A. A., Zhang, C., Xie, T., Tran, L., Dobrin, R., et al. (2013), “Integrated systems approach identifies genetic nodes and networks in late-onset Alzheimer’s disease,” Cell, 153, 707–720.

Zhang, B. and Horvath, S. (2005), “A General Framework for Weighted Gene Co-Expression Network Analysis,” Statistical Applications in Genetics and Molecular Biology, 4, 1–45.

Zhong, Y. and Liu, Z. (2012), “Gene expression deconvolution in linear space,” Nature methods, 9, 8–9.

